# APOBEC3A, not APOBEC3B, drives deaminase mutagenesis in human gastric epithelium

**DOI:** 10.1101/2024.10.28.620744

**Authors:** Yohan An, Ji-Hyun Lee, Joonoh Lim, Jeonghwan Youk, Seongyeol Park, Ji-Hyung Park, Kijong Yi, Taewoo Kim, Chang Hyun Nam, Won Hee Lee, Soo A Oh, Yoo Jin Bae, Junehwak Lee, Jung Woo Park, Jie-Hyun Kim, Hyunki Kim, Hugo Snippert, Bon-Kyoung Koo, Young Seok Ju

## Abstract

Cancer genomes frequently carry APOBEC (apolipoprotein B mRNA editing catalytic polypeptide-like)-associated DNA mutations, suggesting APOBEC enzymes as innate mutagens during cancer initiation and evolution. However, the pure mutagenic impacts of the specific enzymes among this family that are responsible for APOBEC-associated mutagenesis remain unclear in human normal cell lineage, particularly the comparative mutagenic activities of *APOBEC3A* and *APOBEC3B*. Here, we investigated the mutagenic contributions of these enzymes through whole-genome sequencing of human normal gastric organoid lines carrying doxycycline-inducible *APOBEC3A* or *APOBEC3B* cassettes. Our findings demonstrated that transcriptional *APOBEC3A* upregulation led to the acquisition of a massive number of genomic mutations in a few cell cycles. By contrast, *APOBEC3B* upregulation did not generate a substantial number of mutations in gastric epithelium. *APOBEC3B*-associated mutagenesis remained insignificant even after a combined inactivation of *TP53*. Based on the spectrum of acquired mutations after *APOBEC3A* upregulation, we further analyzed *APOBEC3A*-associated mutational signatures, encompassing indels mainly composed of 1bp deletions, characteristics of clustered mutations, and selective pressures operative on cells carrying the mutations. Our observations provide a clear foundation for understanding the mutational impact of APOBEC enzymes in human cells.

## INTRODUCTION

Large-scale cancer genome studies have revealed various mutational processes in human somatic cells (Alexandrov et al. 2020). According to the COSMIC database (Sondka et al. 2024), APOBEC-associated mutations are detected in ∼75% of human cancer types, including bladder transitional cell carcinoma (98%, 381 of 389 samples), breast adenocarcinoma (83%; 759 of 915 samples), and stomach adenocarcinoma (21%; 101 of 486 samples). In addition, APOBEC-associated mutational activity is also observed in non-neoplastic normal cells, particularly within the epithelium of the bladder, bronchus, and small intestine (Lawson et al. 2020; Wang et al. 2023; Yoshida et al. 2020).

APOBEC enzymes are well known for their role in innate immunity, where they destabilize pathogens by inducing DNA and RNA deamination (Vieira and Soares 2013). Recently, they have gained wide recognition as mutagens that also affect host genomes in breast cancer cells (Nik-Zainal et al. 2012b). APOBEC-associated DNA mutations are characterized by predominantly C>T and C>G base substitutions enriched in Tp<u>C</u>pN context (with the mutated cytosine is underlined; Nik-Zainal et al. 2012a). The mutational spectrum has been deconvoluted by two mutational signatures (SBS2 and SBS13), which are consistent across different cancer types (Alexandrov et al. 2020). From the characteristic sequence-context dependency of the mutations, their mutagenic origin was associated with APOBEC enzyme activity (Nik-Zainal et al. 2012b; Lawrence et al. 2013; Roberts et al. 2013). Of the 11 APOBEC family genes in the human genome, *APOBEC3A* (**A3A**) and *APOBEC3B* (**A3B**) have been suggested as major potential contributors to the mutations attributable to SBS2 and SBS13 in most human cell types (Chan et al. 2015; Roberts et al. 2013). In the small intestine, *APOBEC1* was also proposed as a main mutagenic enzyme (Wang et al. 2023).

The mutagenic potential of A3A and A3B has been investigated through a range of human cancer cell lines (Burns et al. 2013; Carpenter et al. 2023; Petljak et al. 2022). APOBEC-associated mutagenic activity was suggested to be episodic in cancer cell line models (Petljak et al. 2019). In addition, non-human model systems, such as *Mus musculus, Saccharomyces cerevisiae,* and *Gallus gallus domesticus*-derived cell lines were also utilized for validating the mutagenic impact of APOBEC enzymes (Durfee et al. 2023; DeWeerd et al. 2022; Naumann et al. 2023; Law et al. 2020; Chan et al. 2015). However, these systems are suboptimal to understand the mutagenicity of APOBEC enzymes in human normal cells, as these models harbor confounding factors, such as additional oncogenic alterations, or non-human genomic backgrounds.

In this regard, we established human gastric organoid lines carrying doxycycline-inducible APOBEC cassettes to evaluate the pure mutagenic activity of A3A and A3B in human normal cells. Combining single-cell cloning and whole-genome sequencing (**WGS**; Jager et al. 2019; Pleguezuelos-Manzano et al. 2020; Youk et al. 2024) as well as duplex DNA sequencing (Hoang et al. 2016), we investigated mutations attributable to A3A and A3B activation in human normal cells.

## RESULTS

### Doxycycline-inducible *APOBEC3A* and *APOBEC3B* in normal gastric organoids

To control the transcription of A3A and A3B in non-neoplastic human cells, we established human gastric organoid lines with doxycycline-inducible expression of either A3A or A3B (referred to as hGO_iA3A_ and hGO_iA3B_, respectively). In these lines, doxycycline treatment simultaneously triggers the upregulation of (1) mCherry fluorescence, and (2) either the A3A or A3B enzyme (**Fig. 1A**). To this end, we utilized *PiggyBac* transposon system (Woodard and Wilson 2015; Wilson et al. 2007) to integrate two independent constructs into the genome of gastric organoids: (1) a cassette expressing rtTA and hygromycin B resistance protein (CMV-rtTA-HygR), and (2) a cassette expressing APOBEC enzymes and fluorescence protein (TRE-APOBEC (A3A or A3B)-IRES-mCherry). Successfully engineered organoids were selected through hygromycin treatment and subsequently constructed into clonal lines. WGS confirmed their clonality (**Supplemental Fig. S1**), copy number and genomic positions of integrated cassettes (**Supplemental Tables S1, S2**), and correctness of the sequence and reading frame of APOBEC gene in the inserted constructs in the hGO_iA3A_ and hGO_iA3B_ lines (**Supplemental Fig. S2**).

**Figure 1.**
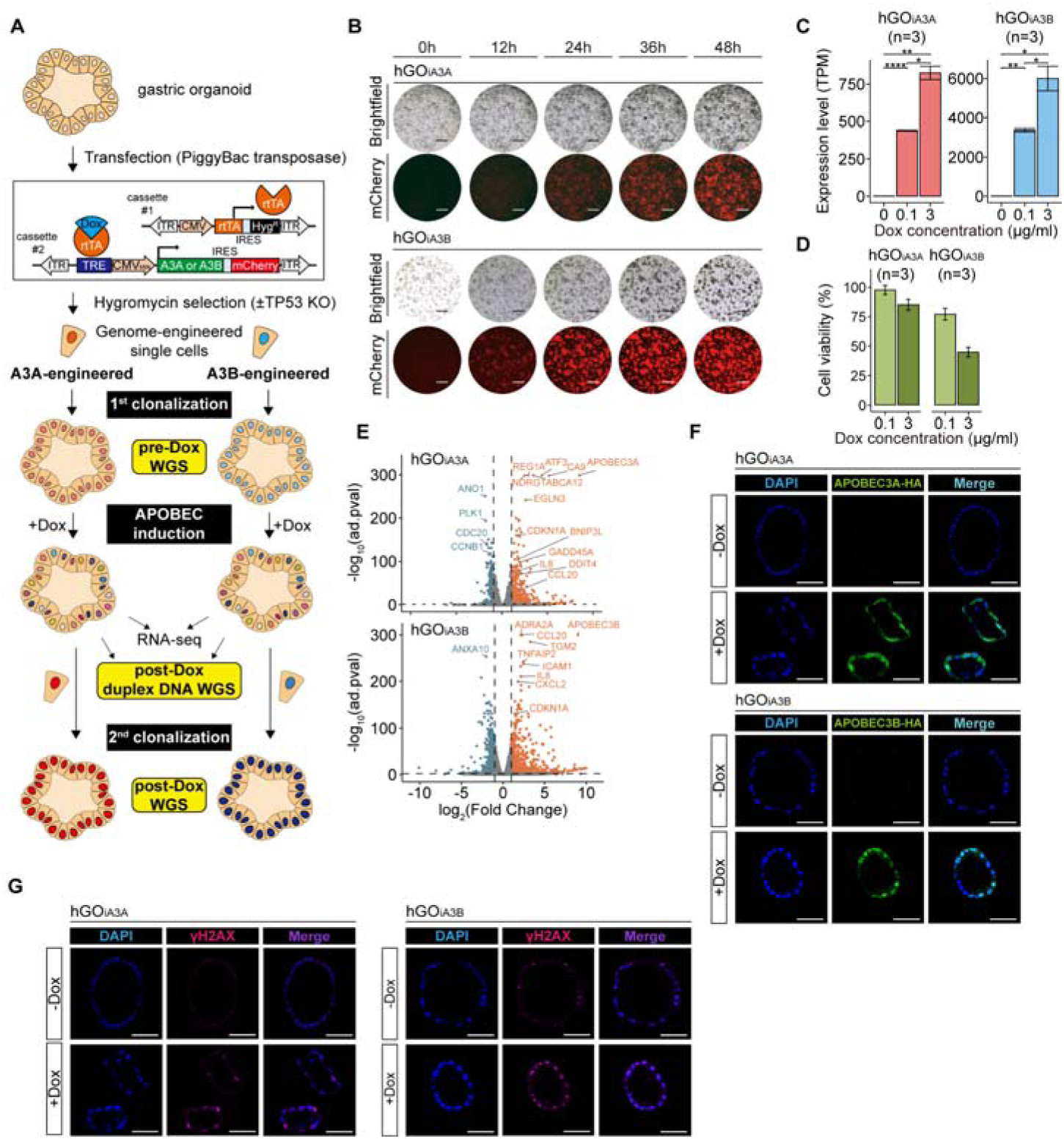
Overview of conditional APOBEC (APOBEC3A or APOBEC3B) overexpression models with human gastric organoids. **(A)** Experimental design of the study. Schematic illustration of genetic engineering, cloning, and sequencing (whole-genome sequencing, duplex sequencing, and bulk RNA-seq) is shown. Dox, doxycycline. **(B)** Changes of morphology and mCherry fluorescence during 48 hours with 0.1 µg/ml doxycycline treatment. (top) hGO_iA3A_ lines and (bottom) hGO_iA3B_ lines. Scale bars represent 1mm. **(C)** Expression levels of APOBEC3A (A3A) or APOBEC3B (A3B) in each line following 0.1 µg/ml and 3 µg/ml doxycycline treatment for 48 hours. (left) hGO_iA3A_ lines (n=3 in each condition) and (right) hGO_iA3B_ lines (n=3 in each condition). Data are presented as mean ± 95% confidence intervals. Statistical significance was determined using a t-test: *p < 0.05, **p < 0.005, ***p < 0.0005, ****p < 0.00005. **(D)** Changes of viability of each line following 0.1 µg/ml and 3 µg/ml doxycycline treatment for 48 hours. (left) hGO_iA3A_ lines (n=3 in each condition) and (right) hGO_iA3B_ lines (n=3 in each condition). Data are presented as mean ± 95% confidence intervals. **(E)** Differentially expressed genes upon APOBEC (A3A or A3B) overexpression following 0.1 µg/ml doxycycline treatment for 48 hours. **(F)** Subcellular localization of overexpressed A3A and A3B following 3 µg/ml doxycycline treatment for 48 hours in each line. (top) hGO_iA3A_ line and (bottom) hGO_iA3B_ line. Scale bars represent 50µm. **(G)** γ-H2AX foci following 3 µg/ml doxycycline treatment for 48 hours in each line. (left) hGO_iA3A_ line and (right) hGO_iA3B_ line. Scale bars represent 50µm.

Upon doxycycline treatment, upregulation of mCherry fluorescence and APOBEC transcripts were clearly observed within 48 hours (**Fig. 1B**). Here, we treated 0.1 µg/ml and 3 µg/ml of doxycycline for low- and high-level induction, respectively. Bulk RNA-seq analysis revealed that gene expression levels of A3A or A3B ranged from 400 to 6,000 transcripts per millions (**TPMs**), representing a several hundred-fold increase compared to background levels (**Fig. 1C; Supplemental Fig.S3**). Of note, transcripts from the endogenous *A3A* and *A3B* genes accounted for only ∼0.2% of total A3A or A3B expression levels following induction, indicating minimal contribution of the innate APOBEC gene copies (**Supplemental Table S3**). The range of expression levels was comparable to the transcriptional levels in single cancer cells with APOBEC expression bursts in many cancer types, including lung adenocarcinoma, head-and-neck squamous cell carcinoma, triple negative breast cancer, esophageal adeno- and squamous cell carcinomas (**Supplemental Fig. S4;** Karaayvaz et al. 2018; Maynard et al. 2020; Puram et al. 2017; Wu et al. 2018). Unfortunately, the levels in gastric adenocarcinoma cells are uncertain, due to the lack of public single-cell transcriptomes through Smart-seq in the cancer type.

Following APOBEC induction, the cell viability of the lines was substantially compromised, suggesting a negative impact of APOBEC enzymes on cell survival (**Fig. 1D**). Differentially expressed genes after APOBEC induction included genes related to cell cycle (e.g., *CDKN1A* and *CDC20*) and immune response (e.g., *IL8* and *CCL20*) in both the hGO_iA3A_ and hGO_iA3B_ models (**Fig. 1E**; Cazzalini et al. 2010; Amador et al. 2007; Matsushima et al. 2022; Wu et al. 2007).

We also observed the nuclear localization of A3A and A3B enzymes following protein translation (**Fig. 1F; Supplemental Fig. S5**). The increased number of γ-H2AX foci after APOBEC upregulation confirmed that the APOBEC enzymes led to DNA damage, such as replication stall or double-strand breaks (**DSBs**) as previously observed (**Fig. 1G; Supplemental Fig. S5**; Burns et al. 2013; Green et al. 2016).

### APOBEC3A, not APOBEC3B, induces genomic mutations

To assess the mutagenic impacts of A3A and A3B, we investigated acquired mutations after 48 hours from doxycycline treatments in the hGO_iA3A_ and hGO_iA3B_ lines. Interestingly, a substantial number of base substitutions, whose spectrum is concordant with a linear combination of SBS2 and SBS13, were detected in the hGO_iA3A_ clones (**Fig. 2A, F; Supplemental Tables S4, S5**). On average, the hGO_iA3A_ lines with 0.1 µg/ml and 3 µg/ml doxycycline exhibited 267 and 2,448 base substitutions genome-wide, respectively (0.1 µg/ml: 95% CI, 107-505; 3 µg/ml: 95% CI, 1,052-3,708; **Fig. 2A, B; Supplemental Table S5**). By contrast, the hGO_iA3B_ clones showed a negligible number of APOBEC-associated mutations (0.1 µg/ml: 95% CI, 0-1; **Fig. 2A, D; Supplemental Table S5**).

**Figure 2.**
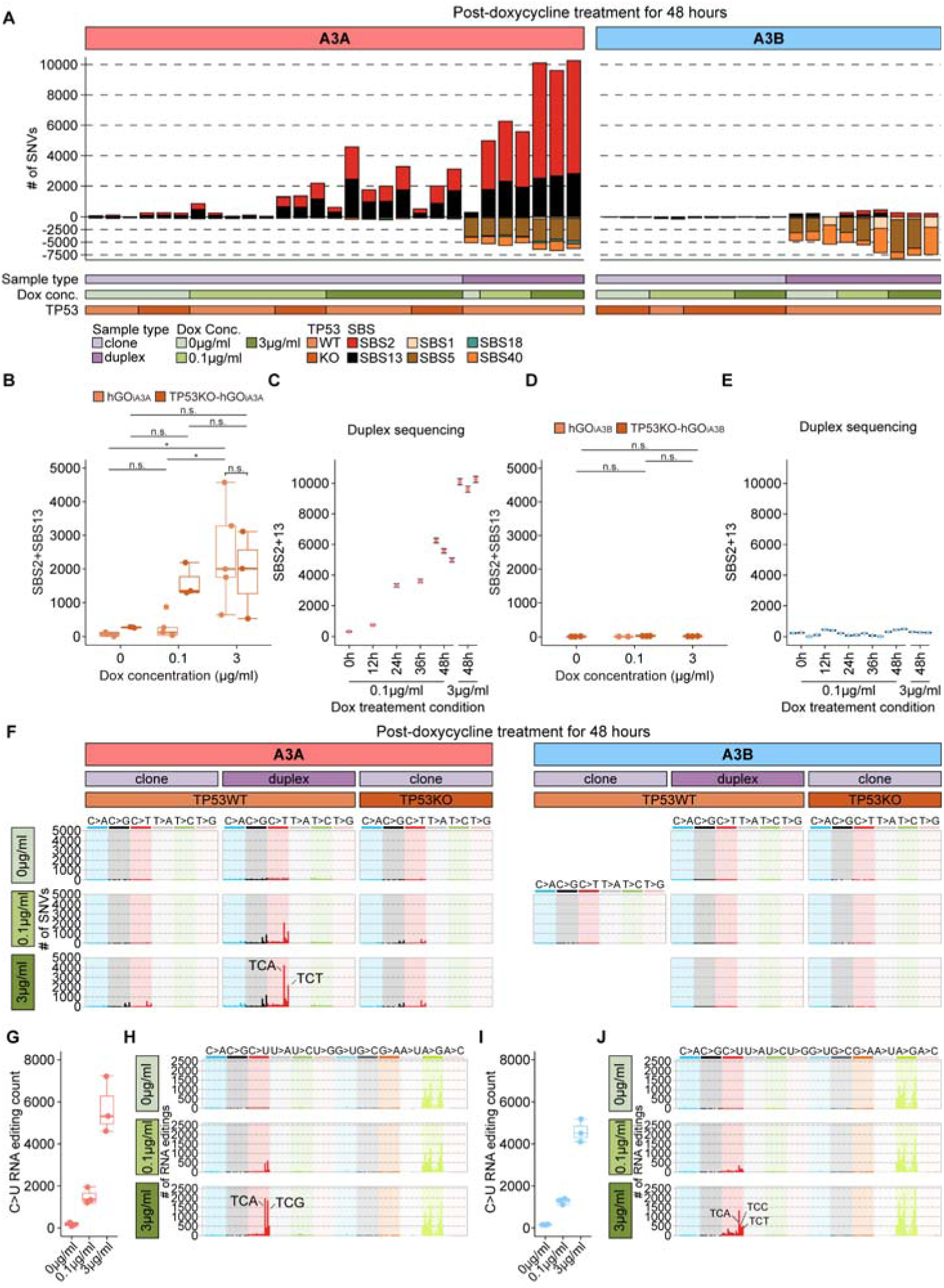
Mutational impact of APOBEC-associated single base substitutions in DNA and RNA. **(A)** Mutational burdens of single nucleotide variants (SNVs) after overexpression of APOBEC (A3A or A3B) measured by whole-genome sequencing of the clones and duplex DNA sequencing classified. (left) hGO_iA3A_ linesand (right) hGO_iA3B_ lines. **(B)** Mutational burdens of APOBEC-associated SNVs in the hGO_iA3A_ and TP53KO-hGO_iA3A_ clone sequencing. The number of A3A-associated SNVs (SBS2+SBS13) in hGO_iA3A_ and TP53KO-hGO_iA3A_ clones under each condition. Statistical significance was determined using a t-test: *p < 0.05, **p < 0.005, ***p < 0.0005,****p < 0.00005. **(C)** The corrected number of A3A-associated SNVs (SBS2+SBS13) in BotSeqS results for hGO_iA3A_ lines under each doxycycline treatment condition. Black lines indicate confidence intervals calculated with *Poisson* distribution. **(D)** Mutational burdens of APOBEC-associated SNVs in the hGO_iA3B_ and TP53KO-hGO_iA3B_ clone sequencing. The number of A3A-associated SNVs (SBS2+SBS13) in hGO_iA3B_ and TP53KO-hGO_iA3B_ clones under each condition. Statistical significance was determined using a t-test: *p < 0.05, **p < 0.005, ***p < 0.0005,****p < 0.00005. **(E)** The corrected number of A3B-associated SNVs (SBS2+SBS13) in BotSeqS results for hGO_iA3B_ lines under each doxycycline treatment condition. Black lines indicate confidence intervals calculated with *Poisson* distribution. **(F)** Mutational burdens and spectra of APOBEC-associated SNVs in each experimental condition. (left) hGO_iA3A_ lines and (right) hGO_iA3B_ lines. **(G)** The number of C>U RNA editing counts in bulk RNA-seq in hGO_iA3A_ lines. n=3 in each condition. **(H)** The spectra of RNA editing in trinucleotide contexts in hGO_iA3A_ lines **(I)** The number of C>U RNA editing counts in bulk RNA-seq in hGO_iA3A_ lines. n=3 in each condition. **(J)** The spectra of RNA editing in trinucleotide contexts in hGO_iA3B_ lines.

Then, we assessed the potential for detection bias in our single-cell cloning system, which could underestimate the mutational impact of APOBEC enzymes. Here, if heavily damaged cells exhibit reduced proliferative potential, they may fail to expand into clones and thus be excluded from subsequent genome sequencing. To bypass the cloning step and directly detect mutations, we employed duplex DNA sequencing technique, enabling the capture of subclonal mutations across the entire cell population, including those present in single cells that could not proliferate further (Hoang et al. 2016).

After 48 hours of doxycycline treatment, the duplex DNA sequencing technique revealed a higher mutational burden of APOBEC-associated base substitutions in hGO_iA3A_ (0.1 µg/ml: 95% CI, 4,590-6,038; 3 µg/ml: 95% CI, 9,316-10,077; **Fig. 2A**; **Supplemental Table S6**). We observed a time-dependent increase in A3A-associated mutational burden following doxycycline treatment (**Fig. 2C**). Our observation confirms that (1) A3A is a potent mutator enzyme, and (2) its quantitative mutational impact may be underestimated in clonal analyses. In contrast, APOBEC-associated base substitutions remained minimal in cell populations exposed to A3B (0.1 µg/ml: 95% CI, 136-352; 3 µg/ml: 95% CI, 79-133; **Fig. 2A**; **Supplemental Table S6**). Of note, we did not detect A3B-associated mutations even at earlier time points, likely prior to cellular clearance (**Fig 2E**), suggesting that the absence of mutations in hGO_iA3B_ is unlikely to be due to negative selection against hypermutated cells.

Despite the minimal mutational impact observed in the organoids, the A3B enzyme was fully functional, exhibiting strong cytosine deamination activity *in vitro*. First, cell lysates from hGO_iA3B_ induced a substantial number of C>T artifacts on extracellular double-strand DNA fragments, particularly in the 5’ head region (∼50bp), with a clear positive correlation to the level of doxycycline concentration (**Supplemental Fig. S6**). As previously reported (Abascal et al. 2021), such artifacts likely arise at single-stranded overhangs of DNA fragments during cell lysis or early stages of library preparation, driven by APOBEC activity in the lysates. Second, recombinant A3B enzymes synthesized from the A3B sequence introduced C>T mutations in cell-free denatured genomic DNA fragments, with nearly all (∼99%) unmethylated (non-CpG) cytosines converted to thymines in approximately 60% of the fragments likely exposed to A3B (**Fig. 3A, B**). In contrast, the remaining ∼40% of the fragments did not show any C>T conversion, suggesting these were not engaged by A3B enzymes. Finally, the increase of γ-H2AX foci in the hGO_iA3B_ following doxycycline induction (**Fig. 1G; Supplemental Figure S5**) also indicates DNA damage consistent with A3B activity. Collectively, our findings suggest that while A3B is enzymatically active, gastric organoids efficiently repair A3B-induced lesions or possess intrinsic mechanisms that suppress A3B-associated mutation in the nucleus, thus limiting its mutagenic impact in the cellular DNA.

**Figure 3.**
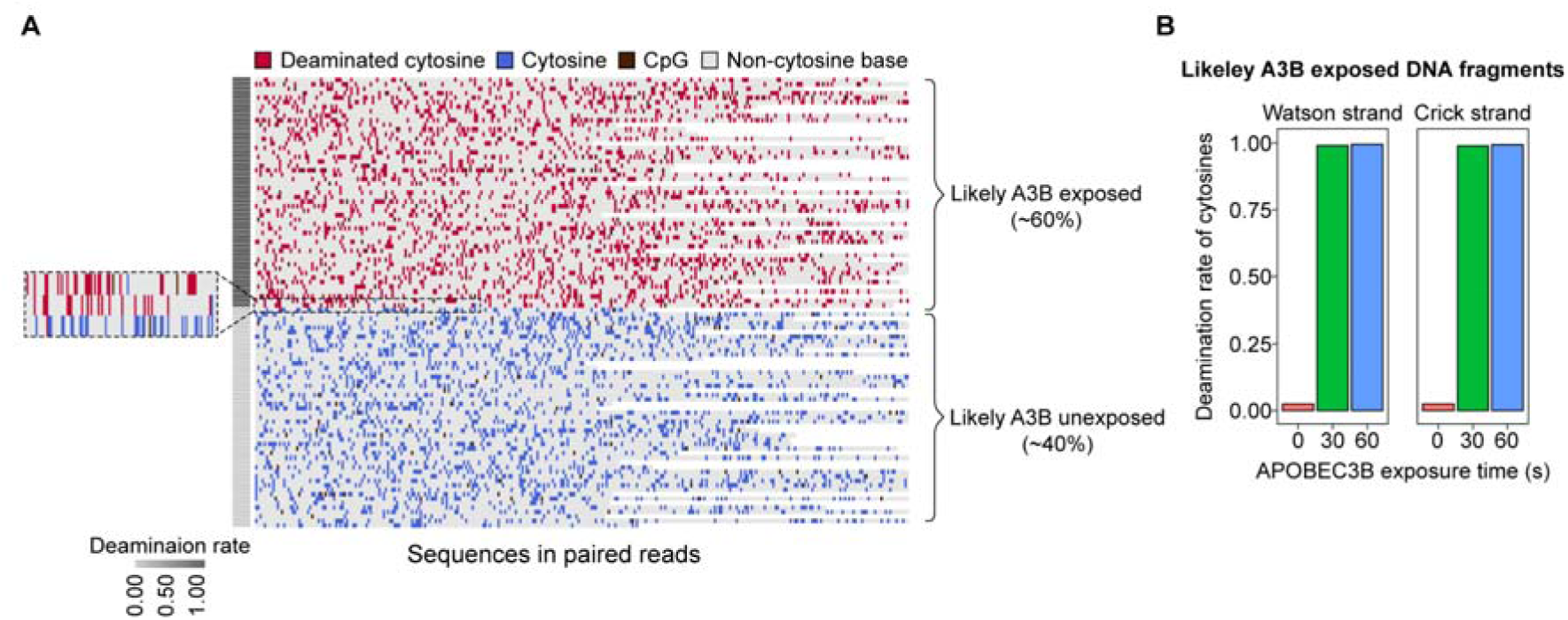
Deamination activity of APOBEC3B *in vitro*. **(A)** Sequences in randomly selected 100 read pairs in APOBEC3B exposed cell-free denatured genomic single stranded DNA for 60 seconds. **(B)** Deamination rate of unmethylated cytosines in A3B-exposed cell-free DNA fragments. CTRL, negative control, 30s, DNA exposed for 30 seconds, 60s, DNA exposed for 60 seconds.

To further investigate whether APOBEC-associated mutagenesis depends on the inactivation of tumor suppressor genes, we generated TP53-knockout derivatives of the hGO_iA3A_ and hGO_iA3B_ lines (hereafter referred to as TP53KO-hGO_iA3A_ and TP53KO-hGO_iA3B_, respectively; **Fig. 1A**). From the TP53KO-hGO_iA3A_ lines, we observed a similar burden of APOBEC-associated mutations to what was observed in the hGO_iA3A_ lines (0.1 µg/ml: 95% CI, 785-1,917; 3 µg/ml: 95% CI, 152-3,091; **Fig. 2A, B; Supplemental Table S5**). Similarly, the TP53KO-hGO_iA3B_ lines also showed results consistent with the hGO_iA3B_ lines, with a lack of APOBEC-associated mutations (0.1 µg/ml: 95% CI, 11-19; 3 µg/ml: 95% CI, 2-12; **Fig. 2A, D**; **Supplemental Table S5**). Taken together, these findings imply that the inactivation of TP53 does not promote APOBEC-associated mutagenesis in gastric organoids.

Despite the increased γ-H2AX foci following A3A or A3B overexpression (**Fig. 1G; Supplemental Figure S5**), which indicates DNA damage and may reflect double-strand breaks in DNA, we did not observe a corresponding increase in structural variations (**SVs**; **Supplemental Table S5**). The absence of APOBEC-induced SVs was also observed in the TP53KO-hGO_iA3A_ and TP53KO-hGO_iA3B_ lines. We speculate that APOBEC-induced DSBs or replication stalls were mostly repaired correctly within the normal cells via the error-free homologous recombination pathway in non-neoplastic gastric cells.

### Both APOBEC3A and APOBEC3B induce C>U RNA editing

Despite their differing mutagenic impact on DNA, both A3A and A3B exhibited deaminase activity on RNA, leading to C>U RNA editing events in the clones. In both the hGO_iA3A_ and hGO_iA3B_ lines, we observed an approximately 8-fold and 30-fold increase in C>U RNA editing following treatment with 0.1 µg/ml and 3 µg/ml doxycycline for 48 hours, respectively, compared to baseline levels (**Fig. 2G, I**). These findings indicate that both constructs possess potent enzymatic activity.

The spectra of C>U RNA-editing deviated from SBS2 for DNA mutations and varied between A3A and A3B (**Fig. 2H, J**). Specifically: (1) both A3A and A3B exhibited reduced specificity for the Up<u>C</u>pU context (equivalent to Tp<u>C</u>pT in DNA); (2) A3A displayed enhanced specificity for the Up<u>C</u>pG context (equivalent to Tp<u>C</u>pG in DNA); and (3) A3B showed decreased specificity for the Up<u>C</u>pN context (equivalent to Tp<u>C</u>pN in DNA). Further *de novo* extraction of RNA editing signatures revealed distinct context-specific characteristics of the A3A and A3B-associated RNA-editing (**Supplemental Fig. S7**), in line with previous reports (Martínez-Ruiz et al. 2023; Fixman et al. 2024; Alonso de la Vega et al. 2023). Expansion of context analysis up to pentanucleotide sequences revealed that A3A-mediated RNA editing were preferentially enriched in ApUp<u>C</u>pApN (equivalent to ApTp<u>C</u>pApN in DNA) and ApUp<u>C</u>pGpN (equivalent to ApTp<u>C</u>pGpN in DNA) contexts (**Supplemental Fig. S8**), whereas A3B-mediated editing did not show such an enrichment.

To further investigate the RNA editing activity of A3A and A3B, we systematically compared the recurrent sites for C>U RNA editing of each enzyme. Among the identified sites, 28,500 were commonly edited by both A3A and A3B, while 558 sites were unique to A3A and 13,078 were specific to A3B (**Supplemental Fig. S9A**). Sequence context analysis revealed that ∼92% of A3A-specific editing events occurred within the Up<u>C</u>pN (equivalent to Tp<u>C</u>pN in DNA) context, compared to ∼69% for A3B-specific sites, suggesting a broader sequence tolerance by A3B (**Supplemental Fig. S9B**). We next examined the local RNA secondary structures at editing sites. Consistent with previous literature (Jalili et al. 2020), A3A preferentially targeted cytosines positioned within defined loop structures, with a clear dependency on both loop size and cytosine position. A3B exhibited similar structural preference; however, the magnitude of these biases was markedly attenuated, suggesting weaker structural constraints on A3B-mediated editing (**Supplemental Fig. S9C**).

### Characteristics of APOBEC-associated mutations

Using the pure A3A-mediated mutational profiles from hGO_iA3A_ lines, we further examined the characteristics of A3A-associated mutational signatures. A3A-associated mutations in the Tp<u>C</u>pA context were 2.7 times more abundant following pyrimidine bases (YpTp<u>C</u>pA) compared to purine bases (RpTp<u>C</u>pA; **Fig. 4A**) as previously reported (Sanchez et al. 2024; Chan et al. 2015). Of note, the RpTpCpA sequence context has been reported as a preferred motif for A3B-associated mutations in the yeast and *in vitro* model (Chan et al. 2015; Sanchez et al. 2024). The YpTpCpA preference in the gastric organoids closely mirrors the enrichment level observed in human cancer tissues carrying APOBEC-induced hypermutations (YpTpCpA/RpTpCpA = 2.5), indicating that A3A could be a key enzyme of the APOBEC-associated hypermutations in most human cancers.

**Figure 4.**
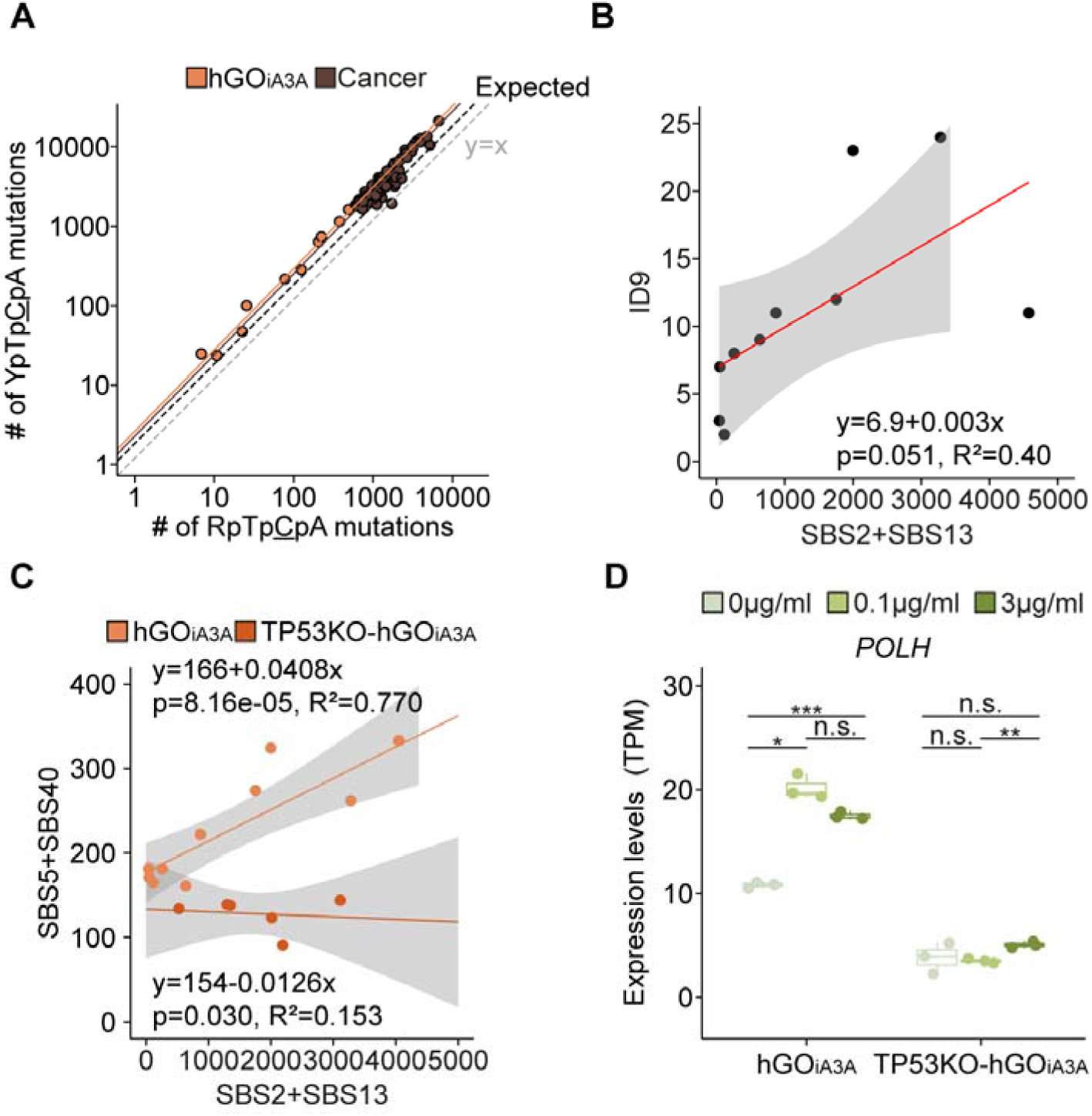
Characteristics of A3A-associated mutational signatures. **(A)** Context preference of A3A between YpTp<u>C</u>pA and RpTp<u>C</u>pA context in hGO_iA3A_ lines and APOBEC-associated mutations in hypermutant cancer samples. Only PCAWG cancer samples with a combined APOBEC-associated mutational burden (SBS2 + SBS13) greater than 5,000 were selected (n=63) among eight cancer types with a high prevalence of APOBEC mutational activity. lung adenocarcinoma (n=15), breast adenocarcinoma (n=12), bladder urothelial carcinoma (n=11), head and neck squamous cell carcinoma (n=13), lung adenocarcinoma (n=6), uterine corpus endometrial carcinoma (n=3), esophageal adenocarcinoma (n=2), and stomach adenocarcinoma (n=1). Dashed black line, expected; orange line, hGO_iA3A_; dark brown line, cancer. **(B)** Correlation between A3A-associated base substitutions and ID9 contributing indels. **(C)** Associations between A3A-associated (SBS2 and SBS13) and age-associated (SBS5 and SBS40) base substitutions among hGO_iA3A_ lines and TP53KO-hGO_iA3A_ lines. **(D)** Gene expression changes of *POLH* (translesion DNA polymerase) after A3A induction in hGO_iA3A_ and TP53KO-hGO_iA3A_ lines.

Further, in line with previous reports from cancer (DeWeerd et al. 2022), indels attributable to the COSMIC ID9 signature (Sondka et al. 2024) showed a suggestive positive correlation with the number of A3A-associated base substitutions in hGO_iA3A_ clones (**Fig. 4B**; p-value = 0.051). The relative rate of ID9-associated indels was one for every 333 SBS2/13 base substitutions in the gastric organoids.

Interestingly, base substitutions attributable clock-like mutational signatures (SBS5 and SBS40) also demonstrated a positive correlation with the burden of A3A-associated base substitutions (SBS2 and SBS13; **Fig. 4C**). This suggests that A3A-induced genomic damage indirectly promotes error-prone DNA repair processes across the genome. However, this association was absent in hGO_iA3A_ clones carrying *TP53* truncating mutations (**Fig. 4C**). We speculate that the difference stems from differential DNA repair pathways according to the activities of TP53 (Kim et al. 2016). Notably, among the polymerases encoded in the human genome, increased transcription of DNA polymerase eta (*POLH*), which is involved in translesion synthesis (TLS; Choi and Pfeifer 2005; Delbos et al. 2005) was exclusively observed in clones without TP53 knockout after induction (**Fig. 4D; Supplemental Fig. S10**). Of note, *REV1*, a component of the translesion synthesis (TLS) pathway, was implicated in the generation of SBS5 and SBS40 (Petljak et al. 2022). These findings collectively suggest that multiple TLS polymerases may contribute to the mutational processes underlying SBS5 and SBS40.

### Clusters of APOBEC-associated mutations

In cancer genomes, APOBEC-induced localized hypermutation events are frequently observed (Nik-Zainal et al. 2012a; The ICGC/TCGA Pan-Cancer Analysis of Whole Genomes Consortium 2020). In our clones, approximately 5% of the 29,650 acquired single-nucleotide variants were clustered within 1 kbp, a frequency 100 times higher than random chance (**Fig. 5A**). Such clustered mutation events can be classified into *omikli* and *kataegis* (usually composed of 2-3 and >3 mutations, respectively) based on statistical models, not by absolute distance (Bergstrom et al. 2022b). A3A-associated clustered mutation events were distributed across 615 *omikli* and 109 *kataegis* events. The observed *kataegis* events ranged from 4 to 22 base substitutions (median = 5). Contrary to the ∼36% of *kataegis* events in cancers reportedly being observed with nearby SVs (The ICGC/TCGA Pan-Cancer Analysis of Whole Genomes Consortium 2020; Bergstrom et al. 2022b; Nik-Zainal et al. 2012a), rearrangement-associated hypermutations were rare in our data (2.8%; 3 out of 109 events). Similarly, the *kataegis* in our clones were independent of other known *kataegis*-shaping genomic events, such as anaplastic DNA bridge (Maciejowski et al. 2020) and extrachromosomal DNA (ecDNA; Bergstrom et al. 2022b). Therefore, our results indicate that the clustered mutations found in our clones were driven by pure activity of A3A.

**Figure 5.**
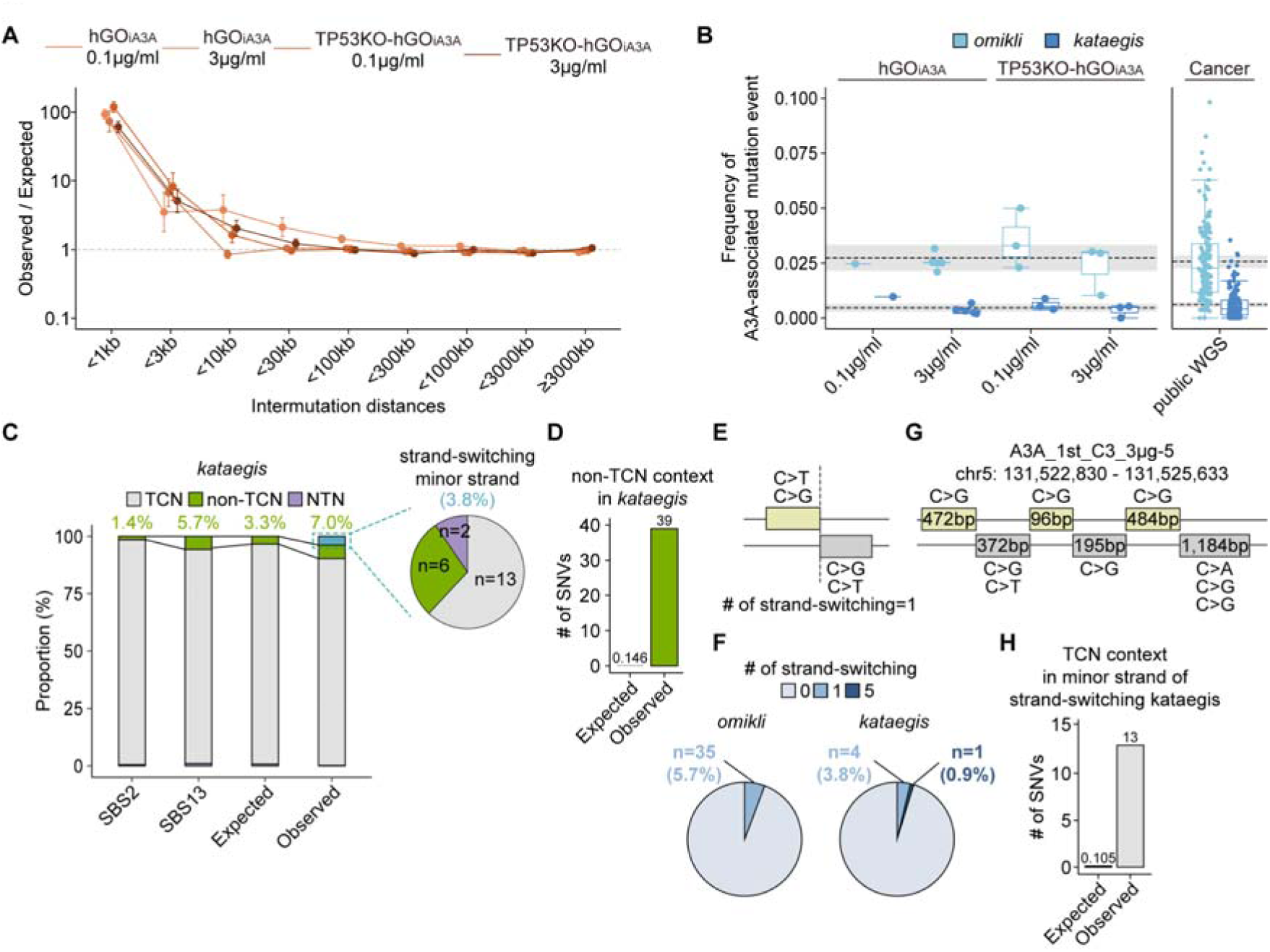
Landscape of APOBEC3A-associated clustered mutation. **(A)** Observed-to-expected ratio of the intermutational distances between substitutions in each clone. Data are presented as mean ± standard error. **(B)** Frequency of clustered mutation events (*omikli* and *kataegis*) among A3A-associated base substitutions in clones and cancer genomes. **(C)** Proportion of classic (in Tp<u>C</u>pN context) and non-classical mutations (in non-Tp<u>C</u>pN and Np<u>T</u>pN) in SBS2, SBS13, and *kataegis* regions in the clones. **(D)** Comparison of expected and observed SNV counts in the non-Tp<u>C</u>pN context within *kataegis* regions. **(E)** Schematic representation of strand-switching *kataegis*. **(F)** Frequency of strand-switching *omikli* and *kataegis*. **(G)** A complex strand-switching *kataegis* with five switches found in clone A3A_1st_C3_3µg-5. **(H)** Enrichments of minor strand mutations in the Tp<u>C</u>pN context in strand-switching *kataegis* events.

The relative frequencies of *omikli* and *kataegis* remained consistent at 2.4% (95% CI: 1.7-3.1%) and 0.4% (95% CI: 0.22-0.58%) of all A3A-associated mutation events, respectively, with each isolated and clustered mutations counted as a single event (**Fig. 5B**; **Supplemental Table S7**). These ratios were more or less constant in clones, regardless of the A3A expression levels and *TP53* mutational status. Interestingly, the frequencies of *omikli* and *kataegis* were comparable to the SV-unrelated *omikli* and *kataegis* frequencies in cancers (**Fig. 5B**; **Supplemental Table S7**).

Within *kataegis* regions, cytosine alterations did not always occur in the Tp<u>C</u>pN context. We observed that ∼7% of cytosine substitutions occurred in non-Tp<u>C</u>pN context (39 out of 556 mutations; **Fig. 5C**), approximately 2-fold higher than expectation based on SBS2 and SBS13 signatures (7.00% vs. 3.33%; chi-square test, p<0.005). Our data indicated these non-Tp<u>C</u>pN mutations arose simultaneously with the *kataegis* for two reasons: (1) Non-Tp<u>C</u>pN mutations were 267-fold more abundant than observed in the background regions (**Fig. 5D**), making contamination by *kataegis*-unrelated mutations unlikely; and (2) Non-Tp<u>C</u>pN mutations within *kataegis* regions were completely phased with other classical Tp<u>C</u>pN mutations on the same chromosome (27 out of 27), where 50% was expected for random contaminating mutations. This suggests that DNA repair mechanisms for cytosine deamination in non-Tp<u>C</u>pN contexts are less accurate during the clustered mutagenesis compared to isolated deamination events.

Of the 615 *omikli* and 106 *kataegis* events (excluding 3 *kataegis* events associated with rearrangements), approximately 5% (35 *omikli* and 5 *kataegis* events) exhibited strand-switching of the mutated cytosines between parental and daughter strands during replication (**Fig. 5E, F**). In one *kataegis* event composed of 9 base substitutions (in clone A3A_1st_C3_3µg-5), we observed five strand-switching events (**Fig. 5G**). Phasing analysis suggested that clustered mutations across both DNA strands originated from single mutation events (26 out of all 26 informative events). Additionally, the mutational spectrum for the mutations in the minorly contributing strand was predominantly composed of mainly C>T or C>G mutations in the Tp<u>C</u>pN context (62%; 13 out of 21 mutations; **Fig. 5C**), implying APOBEC-associated mutations. Besides, the rate of Tp<u>C</u>pN mutations in the minor strand within strand-switching *kataegis* regions was 124-fold higher compared to outside of the strand-switching *kataegis* region (**Fig. 5H**). Although the underlying mechanism is unclear, strand-switching of the A3A enzyme should be possible when generating clustered mutations during replication.

### Epigenetic contexts associated with A3A-associated mutations

Mutational processes are often influenced by the epigenetic contexts of the genome (Otlu et al. 2023). Of the 14 genomic features examined, three features (replication timing, local transcription level, and H3K27me3) showed potential associations with local A3A-associated mutational burdens (**Fig. 6A**). Of the three features, the replication timing and local gene transcription level demonstrated consistent trends correlating with A3A-associated mutation rates (**Supplemental Table S8**).

**Figure 6.**
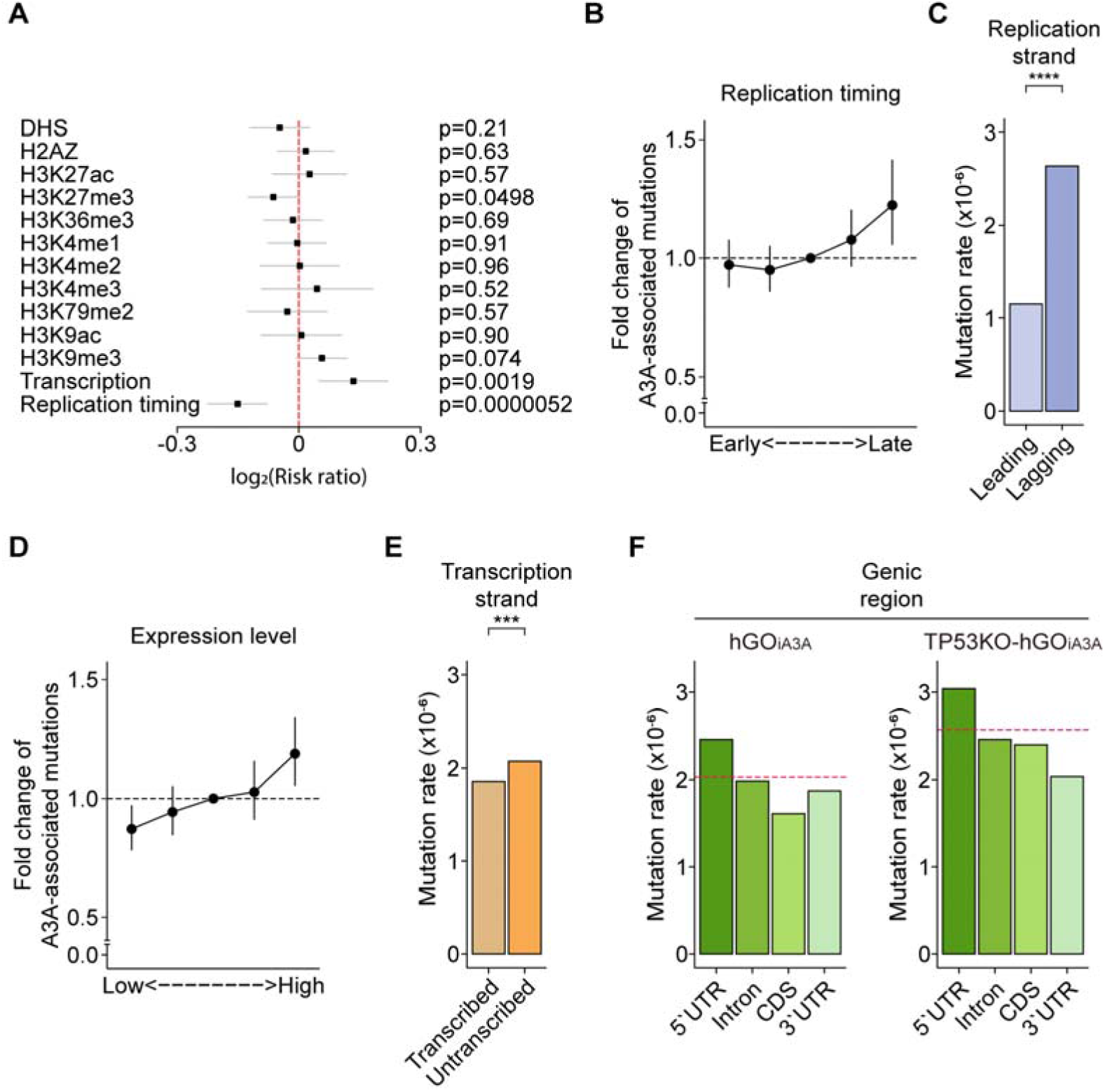
Genomic and epigenomic distribution of APOBEC3A-associated mutations. **(A)** Correlation between epigenetic markers and A3A-associated substitutions. **(B)** Fold change of mutation rates of A3A-associated SNVs according to the replication timing of the genomic region. Data are presented as mean ± 95% confidence intervals. **(C)** Mutation rates depending on the DNA strands during replication (leading and lagging strands). Statistical significance was determined using chi-square test: ****p<0.00005. **(D)** Fold change of mutation rates of A3A-associated SNVs according to the local transcription levels in the genic region. Data are presented as mean ± 95% confidence intervals. **(E)** Mutation rates depending on the DNA strands during transcription (transcribed and untranscribed strands). Statistical significance was determined using a chi-square test: ***p<0.0005. **(F)** Mutation rates within sub-genic regions (5’UTR, intron, protein coding sequences (CDS), and 3’UTR) in (left) hGO_iA3A_ clones and (right) TP53KO-hGO_iA3A_ clones. Red dashed line, average genome-wide mutation rate.

For the replication timing, the latest-replicating regions showed a 1.26-fold higher rate of A3A-associated mutation compared to the earliest-replicating regions (**Fig. 6B**) as previously reported (Kazanov et al. 2015). This may be attributed to the DNA repair mechanisms, including base excision repair, which are particularly active in early-replicating regions consisting of open chromatin (Amouroux et al. 2010; Rhind and Gilbert 2013). Further, the A3A-associated mutation rate in the lagging strand of DNA replication was 1.26-fold higher compared to the leading strand (**Fig. 6C**), which presumably originated from more frequent exposure of single-stranded DNA (ssDNA) induced by Okazaki fragments in the lagging strand (Wu et al. 2020), similar with previous elucidation (Hoopes et al. 2016).

A similar pattern was observed in gene transcription. Mutation rates in genic regions were correlated with expression levels (**Fig. 6D**), showing a 1.37-fold higher mutation rate in actively transcribed genomic regions compared to silent genic regions consistent with the previous study (Kazanov et al. 2015; Nordentoft et al. 2014). Of note, the lagging strand and highly transcribed genes tend to be more frequently single-stranded than the leading strand and silent genes (Okazaki et al. 1968; Gnatt et al. 2001), which may make them more susceptible to A3A-induced DNA damage, respectively. The results demonstrated that DNA regions with frequent ssDNA exposure have a higher chance of being damaged by A3A. In line with the observation, the non-transcribed genic strand was mutated 1.13-fold more frequently than the transcribed strand (**Fig. 6E**), consistent with the previous report (Saini et al. 2017).

Compared with non-coding sequences, protein-coding sequences showed much lower mutation rates, at 0.79-fold the genome average, suggesting a selective pressure against mutations that could alter amino-acid-changing mutations (**Fig. 6F**). Interestingly, in TP53-inactivated clones (TP53KO-hGO_iA3A_ clones), mutation rates in protein-coding regions increased to 0.834-fold of the genome average, potentially due to reduced negative selection pressures in the absence of functional *TP53*.

## DISCUSSION

Our study clearly demonstrated the qualitative and quantitative mutational impact of A3A and A3B in human non-neoplastic cells using a gastric organoid culture system. The gastric organoid model was particularly suitable for this study based on three key criteria: (1) the biological relevance, supported by the presence of significant APOBEC-associated mutations in corresponding gastric cancer tissues, (2) the robust proliferative capacity under culture conditions, enabling experimental feasibility, and (3) the availability of standardized protocols for genetic manipulation. Although APOBEC-associated mutations are also frequently observed in breast and lung cancers, we did not include normal breast or lung epithelial cell models for this study due to their limitations in the cell proliferation capacity in organoid culture condition and genetic engineering compatibility.

When A3A was induced by 3 µg/ml of doxycycline in hGO_iA3A_ lines, transcription was activated for ∼2-3 days, reaching peak gene expression levels of ∼800 TPM. Under these conditions, we detected ∼2,500 A3A-induced base substitutions in the proliferative clones, suggesting that ∼1,000 base substitutions could be acquired in a day. As demonstrated by duplex sequencing, the mutational burden is likely even higher in less proliferative cells.

These findings support the notion that a transcriptional burst of A3A can also lead to a massive number of mutations in normal gastric epithelium, consistent with previous report in cancer (Petljak et al. 2019).

Our models indicated increased mutational burdens of SBS5 and SBS40 in proportion to the overall burden of A3A-associated mutations. These signatures were significantly enriched in late-replicating regions and correlated with multiple epigenomic features, including replication timing and transcriptional activity, consistent with previous reports (**Supplemental Fig. S11;** Sondka et al. 2024). Together, these findings support a model in which A3A-induced DNA damage perturbs global DNA repair processes, thereby promoting the accumulation of SBS5 and SBS40 mutational signatures.

Previous studies have highlighted the mutagenic potential of A3B across various model systems (Dananberg et al. 2024; Carpenter et al. 2023; Durfee et al. 2023; Chan et al. 2015). In contrast, a recent study reported only a modest reduction in APOBEC-associated mutations after A3B inactivation in human cancer cell lines, suggesting a more limited role for A3B in mutagenesis (Petljak et al. 2022). To our knowledge, our study is the first to directly assess the mutagenic activity of A3B in human normal cells. Our findings indicate that A3B alone does not act as a major mutator, at least in the context of human normal gastric epithelium.

Understanding the potential cell-type specificity of the mutagenic activity of A3B, as well as the underlying mechanisms, represents an important direction for future research. Applying similar experimental approaches to organoids derived from a wider range of tissue types, in both normal and neoplastic contexts, will be essential to clarify the broader relevance of A3B activity. Furthermore, emerging technologies that enable simultaneous whole-genome and transcriptome sequencing at the single-cell level will be invaluable for resolving the timing, cellular context, and consequences of APOBEC-associated mutagenesis in a tissue- and model-independent manner.

Although this study successfully evaluated the mutagenic activity of A3A and A3B in human gastric epithelium, it has a few technical limitations that warrant consideration. First, the endogenous copies of A3A and A3B were not inactivated in the hGO_iA3A_ and hGO_iA3B_ models, raising the possibility that a small fraction of the observed APOBEC-associated mutations may have originated from these native enzymes. In addition, the inclusion of further control models, including catalytically inactive mutants of A3A and A3B (i.e., A3A-E72A and A3B-E255A; Carpenter et al. 2023), would help more definitively rule out the contribution of indirect mechanisms triggered by APOBEC overexpression. In parallel, treatment with APOBEC-enzyme inhibitors to our model could also help to validate A3A as a directly mutagenic enzyme. However, we consider these experiments to be of limited necessity, as (1) the exogenously introduced APOBEC enzymes is quantitatively predominant in our system (**Supplemental Table S3**), and (2) it is widely accepted that APOBEC enzymes are direct mutators with intrinsic mutagenic activity. Lastly, though our gastric organoid models reproduced comparable transcriptional bursts of APOBEC enzymes *in vivo*, they cannot fully replicate the physiological complexity of *in vivo* systems.

## METHODS

### Materials Availability

Organoids established in this study will be available under a material transfer agreement. To do so, please contact the lead author.

### Human normal gastric samples

Normal gastric tissues were obtained via endoscopic biopsy from a female undergoing routine screening. The protocol for this study was approved by the institutional review board of Yonsei University Gangnam Severance Hospital (3-2018-0207) and KAIST (KH2022-211).

### Confirmation of non-neoplasticity of normal gastric organoids with whole-genome sequencing

Whole-genome sequencing analysis confirmed the absence of potent cancer driver mutations and copy number variations of DNA (**Supplemental Fig. S12**) in the normal gastric organoids.

### Human stomach organoid culture

Organoid culture methods and media compositions were adopted from previous research with slight modification (Bartfeld et al. 2015). Wnt3A and R-spondin-1 conditioned media was produced with HEK293 cell line producing Afamin-Wnt3a (Mihara et al. 2016) and Cultrex HA-R-Spondin 1-Fc 293T cell line (Trevigen). Complete medium composed of Wnt3A conditioned medium, R-spondin-1 conditioned medium, Advanced DMEM/F-12 (Gibco, Cat No.12634028), HEPES (1M) (Gibco, Cat No.15630-080), Penicillin/streptomycin (10,000 U/mL) (Gibco, Cat No.15140122), GlutaMax (Gibco, Cat No.35050061), Human EGF Recombinant Protein (Invitrogen, Cat No.PHG0311), hNoggin (Peprotech, Cat No.120-10C), hFGF10 (Peprotech, Cat No.3100-26), B27 supplement (50x) serum free (Gibco, Cat No.17504044), N-acetylcysteine (Sigma-Aldrich, Cat No.A9165), Gastrin (Sigma-Aldrich, Cat No.G9145), Y27632 (Sigma-Aldrich, Cat No.Y0503), TGF-b R kinase inhibitor IV (Biogems, Cat No.3014193; **Supplemental Table S9**).

Tissues were dissociated into 10∼15 clustered cells by pipetting with 1000µl tips after 30 minutes of incubation in 4°C in TrypLE (Gibco, Cat No.12604013). Tissues were washed twice with PBS, and the pellet was isolated by removing the supernatant after centrifugation at 300g for 5 minutes at 4°C. The resuspended pellets with cold Matrigel (Corning, Cat No.BDL356231) were seeded on the 12- or 24-well plates (Merck) in dorm shape. After incubation of plates in the humidified incubator (5% CO2, 37) for 10 minutes to polymerize Matrigel, prewarmed 0.5-1 ml of culture medium was added to the well. Organoids were cultured in the humidified incubator. The medium was changed every two to three days.

Organoid passaging was conducted for about two weeks. Medium in the well was replaced by cell recovery solution (Corning, Cat No.354253), and the plates were incubated for 40∼60 minutes at 4°C to dissolve Matrigel. Organoid pellet was isolated by removing supernatants after centrifugation at 300g for 5 min at 4°C. Organoids followed by incubation for 5 minutes at 37°C with resuspension with Accutase (Stemcell Technology, Cat No.07922) were dissociated into 10-15 cell clusters by pipetting 10-20 times. After diluting Accutase by adding 1 ml of ADF media (Gibco), the pellet was isolated by centrifugation at 300g for 5 minutes at 4°C. Organoids were seeded in a 12- or 24-well plate at a ratio of 1:4 to 1:6 within Matrigel.

### Preparation of vectors for transfection

Two vector systems: (1) CMV-rtTA-HygR vector (Addgene, Cat No.102423) / and (2) CRISPR-*Cas9* vectors containing gRNA sequence for TP53 (Addgene, Cat No.121917) were purchased from Addgene. Basically, mini-prep and maxi-prep were conducted with commercial competent cells (Biosearch Technologies, Cat No.6010) and mini-prep kit (QIAGEN, Cat No.27104) and maxi-prep kit (QIAGEN, Cat No.12123) according to the manufacturers’ protocols. To create pPB-CMVmin-A3A/A3B-IRES-mCherry vectors, APOBEC constructs were designed following these steps. NCBI reference sequences for *APOBEC3A* (NM_145699.4) and *APOBEC3B* (NM_004900.4) were utilized for the design. Kozak consensus sequences (GCCACC) were added at the 5’ end of the A3A/A3B cDNA sequences, while HA tag sequences (5’-TAC-CCA-TAC-GAT-GTT-CCA-GAT-TAC-GCT-3’) were appended to the 3’ ends of the A3A/B cDNA sequences. Subsequently, *XhoI* and *NotI* restriction enzyme cut sites (CTCGAG and GCGGCCGC) were added to the 5’ and 3’ ends of the construct, respectively. *De novo* gene synthesis from GenScript (Piscataway, NJ) was used for the synthesis of the two constructs. Following the acquisition of the A3A and A3B constructs, the constructs were cloned into the backbone vector (pPB-CMVmin-TRE-IRES-mCherry (Lee et al. 2022) to make pPB-CMVmin-A3A/B-IRES-mCherry cassette.

### Transfection of organoids

Organoids were prepared with the same dissociation protocol of passaging organoids. Transfection methods were adopted from previously reported protocols (Gaebler et al. 2020; Fujii et al. 2015). A mixture of three types of vectors were utilized for transfection: having (1) TRE-APOBEC (A3A or A3B)-IRES-mCherry cassette and (2) CMV-rtTA-HygR cassette (3) pPiggyBac transposase cassette. For hGO_iA3A_ lines, organoids were suspended in 90µl of Opti-MEM (Gibco, Cat No.31985062), and mixed with about 30 µg of each vector. Electroporation was conducted using a previously described program from the literature (Fujii et al. 2015); **Supplemental Table S10**). Transfected organoids were cultured for seven days with the medium composition described in the previously reported protocol. Organoids were incubated with the medium containing 1 µg/ml hygromycin (InvivoGen, Cat No.ant-hg-1) for seven days after splitting. To isolate the transfected organoids having insertions of two vector cassettes for conditional overexpression, organoids were incubated with doxycycline (Sigma-Aldrich., Cat No.D9891-1G) in a 3 µg/ml containing medium for 12-16 hours. mCherry-positive organoids were manually isolated by pipetting under a fluorescent microscope, and single-cell cloning was conducted according to a previously reported method by using FACSAria (BD Biosciences) and manually picking single cell originated organoids by pipetting (Youk et al. 2021). Isolated organoids were dissociated in the same way to passage. Then, organoids were then filtered through a 40 μm strainer (Falcon, Cat No.352340). Using FACSDiva software, pure single cells were isolated. After seeding the organoids, the single-cell-originated organoids were isolated. For hGO_iA3B_ lines, organoids were suspended in 100 µl of BTXpress buffer (BTX), and 10 µg of vector mixtures were mixed. The following steps were carried out in the reported protocol (Fujii et al. 2015). Selection and single-cell cloning steps were conducted using the same method as for hGO_iA3A_ lines.

### Doxycycline treatment

A doxycycline stock solution was prepared by dissolving doxycycline in the DMSO. The stock solution was added to the medium to achieve final concentrations of 0.1 µg/ml and 3 µg/ml. Before treatment, dissociated 10k viable cells were seeded in 24-well the plates. After 7 days, doxycycline solution was added to the medium, and the organoids were incubated with doxycycline.

### Viability assay of organoids

According to the manufacturer’s protocol, the viability of organoids was measured using a Celltiter-Glo 3D Assay kit (Promega, Cat No.G9681). Each well was washed twice with 1 ml of PBS by repeatedly removing the medium and adding PBS. After washing, PBS was replaced with fresh medium. An equivalent volume of reaction solution was added to each well. Matrigel was dissolved in the reaction solution by incubating at 4 for 30 minutes with shaking. The amounts of ATP in each well were measured with a luminometer (BERTHOLD Technologies GmbH, Cat No.TriStar LB 942). The viability percentage was calculated by dividing luminescence after doxycycline treatment by the average luminescence of untreated control groups.

### Capturing fluorescent images

Fluorescents of mCherry after doxycycline treatment were monitored under the fluorescent microscope (LEICA, DMi8; excitation = 541-551 nm; emission = 565-605 nm). Fluorescent images were captured using Las X programs provided by the manufacturer, and brightness/contrast were adjusted using the same Las X program.

### Immunostaining and imaging of organoids

Whole-mount staining of human gastric organoids was performed as previously described (van Ineveld et al. 2020). Briefly, organoids were fixed with 4 % paraformaldehyde (PFA) after depolymerizing the 3D matrix using an ice-cold cell recovery solution (Corning). After washing the cells with 0.1% PBS-Tween-20 and blocking with organoid washing buffer (0.1 % Triton X-100, 0.02 % SDS, 0.2 % bovine serum albumin (BSA) in 1X PBS), immunolabeling was performed with mouse anti-HA tag antibody (Santa Cruz, Cat No.sc-7392; 1:50) and rabbit anti-phospho-histone H2A.X antibody (CST, Cat No.2577S; 1:400) to detect inserted HA-tagged APOBEC proteins and double-strand breaks, respectively. Goat anti-mouse Alexa Fluor 488 (Thermo Fisher Scientific, Cat No.A-11001; 1:500) and donkey anti-rabbit Alexa Fluor 647 (Thermo Fisher Scientific, Cat No.A32795; 1:500) were used as secondary antibodies, and nuclei were counter-stained with DAPI (Sigma-aldrich., Cat No.D9542; 1:1000). After washing the samples, FUnGI clearing solution was added to the organoids. The samples with the clearing solution were directly mounted between two coverslips with a 0.25 mm-deep iSpacer (SunJin Lab, Cat No.IS213), and the imaging was performed at least 1 hour after slide preparation.

Imaging was performed with a Leica Stellaris 8 Confocal Microscope at the Research Solution Center (RSC) at the Institute for Basic Science (IBS). Alexa Fluor 488 (excitation, 479 nm; emission, 482–554 nm), Alexa Fluor 647 (excitation, 580nm; emission, 663-716 nm), and DAPI (excitation, 405 nm; emission, 425-489 nm) signals were obtained. Either 40X or 63X objectives with a digital zoom factor of 1 to 7-fold were used. The X/Y resolution was set to 1024 x 1024 pixels. Images were processed and analyzed using Adobe Photoshop.

### Standard whole genome sequencing and alignment

Organoids were isolated by removing matrigel with a cell recovery solution (Corning), similar to the method used for passaging organoids. According to the manufacturer’s protocol, DNA from clonal organoids was extracted using DNeasy Blood & Tissue Kit (QIAGEN, Cat No.69506). Libraries were constructed with Truseq DNA PCR-Free Library Prep Kits (Illumina, Cat No.20015963) according to the manufacturer’s protocols. Whole genome sequencing was conducted by the NovaSeq 6000. Organoid clones were whole-genome sequenced with a mean 30-fold depth of coverage. Adapter sequence in the FASTA files was removed by Cutadapt software (https://cutadapt.readthedocs.io/en/stable/; Martin 2011). Sequencing reads were aligned to the human reference genome, GRCh37, with bwa-mem v0.7.17 (https://github.com/lh3/bwa; Li and Durbin 2010). Further processing, including sorting, marking duplication, and indel realignment, was conducted with samtools v1.9 (https://www.htslib.org/; Li et al. 2009), PICARD v2.1.0 (https://broadinstitute.github.io/picard/; McKenna et al. 2010), and GATK tools v3.8.0 (https://gatk.broadinstitute.org/hc/en-us/categories/360002369672-Tool-Index; McKenna et al. 2010).

### BotSeqS library construction

DNA for the BotSeqS library was extracted with All-Prep DNA/RNA Mini kit (Qiagen, Cat No.80204) according to the manufacturer’s protocol. Constructed libraries with Truseq DNA PCR-Free Library Prep Kits (Illumina) were utilized for subsequent steps. The construction of BotSeqS libraries was based on the previous literature with slight modifications (Hoang et al. 2016). Briefly, the quantification of DNA libraries was conducted with KAPA Library Quantification Kit ILLUMINA® Platforms (Roche, Cat No.KK48247) according to the manufacturer’s protocol. An effective library concentration was calculated considering the size of added DNA length during library construction. PCR amplification of libraries equivalent to 4 pg of DNA was performed with the KAPA HiFi Uracil+ kit (Roche, Cat No.KK2802) and primers having a Y-adaptor sequence with a phosphorothioate bond(*) at the 3’ end from IDT (Coralville, IA):

Forward: 5’-AATGATACGGCGACCACCGAG*A-3’

Reverse: 5’-CAAGCAGAAGACGGCATACGA*G-3’

PCR programs were adopted from the protocol of KAPA Library Amplification Kits (Roche), and PCR was conducted with 20 cycles. Library clean-up was performed with SPRIselect beads (BECKMAN COULTER life Sciences) according to the manufacturer’s protocol. The libraries were sequenced as paired-end sequencing (2x151 bps) using NovaSeq 6000.

### *In vitro* deamination of DNA with synthesized APOBEC3B

The NEBNext Enzymatic Methyl-seq (EM-seq) Kit (NEB, Cat No.E7120S) and the manufacturer’s protocol were utilized for the construction of the DNA library with slight modification. Firstly, the DNA library was constructed with DNA extracted from a doxycycline treatment-free hGO_iA3B_ line. DNA libraries equivalent to 10 pg of input DNA were prepared following the process used for BotSeqS library construction. Oxidation of DNA by TET2 and denaturation steps were performed according to the manufacturer’s instructions. For deamination steps, synthesized recombinant APOBEC3B from EUROPROTEIN INC. (North Brunswick Township, NJ) was used instead of the APOBEC protein included in the EM-seq kit. Additionally, the reaction time was modified from 3 hours to 30 and 60 seconds. Library amplification steps followed the BotSeqS library construction.

### Whole transcriptome sequencing library construction

RNA was isolated during the extraction of DNA for BotSeqS libraries with AllPrep DNA/RNA Mini kit according to the manufacturer’s instructions. RIN value of raw RNA was measured with Agilent 4200 TapeStation (Agilent, Cat No.G2991BA). Libraries were constructed with NEBNext Ultra II Directional DNA Library Prep Kit for Illumina (NEB, Cat No.E7760) and QiAseq FastSelect -rRNA HMR kit (Qiagen, Cat No.334388) according to manufacturers’ instructions. Clean-up process was conducted with SPRIselect beads according to the manufacturer’s protocol. Libraries were sequenced with Novaseq 6000 with paired-end sequencing.

### Confirming non-neoplasticity in primary human gastric organoid

The germline mutation calling mode in GATK v4.0.12.0 (https://gatk.broadinstitute.org/hc/en-us/categories/360002369672-Tool-Index; McKenna et al. 2010) was utilized for the identification of mutation in the organoid lines. The primary call set was first filtered by in- house scripts using the pysam module in Python (https://pysam.readthedocs.io/en/latest/index.html; Li et al. 2009) with the following conditions: (1) the number of reference reads ≥ 5; (2) the number of variant reads ≥ 3; (3) median miss-matched bases in variant reads < 5; (4) median mismatched bases in variant reads < 5 (5) median mapping quality of reference reads > 20 (6) median mapping quality of variant reads > 20 (7) median distance from the ends of reads > 4. The impact of variants in the protein-coding region was annotated with ANNOVAR (https://annovar.openbioinformatics.org/en/latest/; Wang et al. 2010). Non-silent mutations located in the protein-coding and splicing regions were isolated. The functional impact in cancer driver genes (the list can be downloaded at https://cancer.sanger.ac.uk/cosmic) was evaluated with PolyPhen-2 (Adzhubei et al. 2013) and SIFT (Ng and Henikoff 2003) scores annotated by ANNOVAR, AlphaMissence (https://alphamissense.hegelab.org/; Cheng et al. 2023), and MutPred-Indel (http://mutpred2.mutdb.org/mutpredindel/about.html; Pagel et al. 2019). Considering the role of cancer driver genes, the impact of variants was finally assessed.

### Calling copy number variations

Copy number variations were accessed by Sequenza (https://sequenza-utils.readthedocs.io/en/latest/<u>#;</u> Favero et al. 2015). Copy number variations in segments smaller than 1 Mbp were considered false positives. After removing the copy number variations in short segments, final copy number variations were investigated by a second run of the Sequenza.

### Detection of somatic mutations

To detect single nucleotide variants and indels in clonal organoids, GATK Mutect2 v4.1.9 (https://gatk.broadinstitute.org/hc/en-us/categories/360002369672-Tool-Index; McKenna et al. 2010) and Strelka2 v2.9.2 (https://github.com/Illumina/strelka; Kim et al. 2018) were utilized. Bulk whole-genome sequencing of mother clones, hGO_iA3A_,TP53KO-hGO_iA3A_, hGO_iA3B_, and TP53KO-hGO_iA3B_ organoids, were utilized as matched normal for calling somatic mutations in doxycycline treated organoids. In each call set, false-positive variants were filtered out with in-house python scripts annotating information within BAM files with the pysam, a python module. The filtered call sets from two callers, Mutect2 and Strelka2, were merged, and the union call set was utilized for downstream analysis. To exclude mutations generated during the culture of mother clones, recurrent somatic mutations observed across multiple doxycycline-treated clones, including control samples, were removed from each call set.

In the case of BotSeqS, VarScan2 v2.3.9 (https://varscan.sourceforge.net/; <u>Koboldt et al. 2012)</u> was used to increase the sensitivity of calls. Similarly, in-house Python scripts were utilized to filter out false-positive calls. Specifically, F1R2 and F2R1 reads from same DNA was grouped and filtered with the following criteria: (1) total depth of each type of read ≥ 3; (2) the number of variant reads ≥ 3 or 90% of reads; (3) distances of mutations from each end of each read > 5 bp; (4) distances of mutations from the end of reads > 100 bp considering the DNA strand (e.g., excluding C>T near the end of F1R2 reads, but not G>A); (5) median mapping quality of variant reads ≥ 20; (6) median base quality of variant reads ≥ 30; (7) the number of variants in WGS of HEK293T <3; (8) the number of variants in WGS of control < 1. Unlike with standard whole genome sequencing, C>T mutations were frequently observed at the end of the reads, and these were considered false positives and removed. Additionally, rare variants from HEK293T cells, which were used to generate conditioned medium, were observed. Thus, variants observed in HEK293T cell line BAM files at least three times were removed from the call set.

### Calculation of mutation rate in BotSeqS

Unlike standard WGS, the length of the effective covered region was calculated for each BAM file. First, each read was evaluated using in-house scripts that considered DNA strand orientation and applied the following criteria to isolate effective DNA fragments: (1) median mapping quality > 20 (2) total depth of each type of reads ≥ 3. Only regions where both F1R2 and F2R1 reads were aligned were included in the covered region calculation. To account for the exclusion of mutations located within 100 bp of extreme of read ends during variant filtering (considering the reference genome strand), the total length of the covered region was adjusted by multiplying it by (151x2-110)/(151x2)). Mutation rates were calculated by dividing the number of observed mutations by this adjusted covered length. To normalize mutational burdens against the standard genome length, mutation rates were then multiplied by the total genome length excluding repeat regions (3,041,373,115 bp) and further doubled.

### Detection of deamination during *in vitro* denomination with APOBEC3B

The calling pipeline for BotSeqS results was utilized with slight modifications. Among the criterias, the filtering criteria for distances from the ends of reads were neglected. Initially, all of the filtered mutations were collected. Among the mutations in grouped DNA, only those in DNA fragments where only one strand of DNA was mapped were counted.

To calculate the genome-wide mutation rates, the BotSeqS pipeline was also used, with the exception of counting cytosine or guanine considering the strand of DNA. Since original methylated CpG sites were preserved during the library construction, total counts and mutation counts in all of CpG contexts were excluded. To calculate mutation rates in DNA fragments with at least one C>T variant, ranges of DNA fragments overlapping with the variants were isolated while considering the DNA strands using BEDtools (https://bedtools.readthedocs.io/en/latest/; Quinlan and Hall 2010). After this step, cytosine or guanine outside of CpG contexts were counted.

### Analysis of mutational signatures

Mutational signature analysis of single base substitutions (SBS) and small indels were carried out using non-negative least squares method. The mutational signature was represented by 96 patterns of SBS and 83 patterns of small indels (Alexandrov et al. 2020). Pre-learned catalogs of mutational signatures in the COSMIC (Sondka et al. 2024) were used to fit individual samples with a known set of signatures for each tissue type. SBS2 and SBS13 (known APOBEC signatures) were included for all cases.

### Analysis of mutation rates depending on epigenetic marker

Risk ratio of mutation rates in non-signal regions and signal-detected regions depending on each epigenetic marker was analyzed based on previous researches (Nam et al. 2023; Supek and Lehner 2017). Genome-wide signals of each marker, including replication timing, across eight cell types (E017,E114, E117, E118, E119, E122, E125, and E127) were downloaded from Roadmap Epigenomics Consortium. Fold-enrichment signal was averaged across the cell types, and the regions showing fold-enrichment signal lower than 1 identified as bin 0, non-signal regions. The other regions were classified as the signal-detected regions. Based on the defined regions, the number of SNVs were counted, and subsequently, risk ratio was calculated considering 3bp genomic context. For APOBEC-associated mutations, only cytosines within the TCN context were considered as background, and C>T and C>G substitutions at these sites were counted. For SBS5 and SBS40, thymine bases outside the TCN context were used as the reference, and T>A,T>G,T>C, C>T and C>G substitutions were counted. C>A mutations were excluded to avoid confounding effects from overlapping sequence contexts associated with SBS1 and SBS18.

To analyze fold-enrichment of mutation rates, in case of replication timing and H3K27me3 signals, the signal-detected regions were further divided into four groups, considering the equal distribution of length of each group. In case of expression level, expression level with TPM unit in each base calculated from merged whole-transcriptome sequencing data of doxycycline treated organoids for 48 hours under 3 µg/ml doxycycline was utilized. First of all, non-genic regions and genic regions were classified based on gene information from the “hg19_refGene.txt” file. Only genic regions were included in the analysis. Genic regions having 0 of TPM were defined as bin0 regions. TPM>0 regions were classified into 4 groups (cutoff; bin1=0.05, bin2=1.73, and bin3&bin4=9.68) based on quartiles of TPM. Considering interference among H3K27me3, replication timing, and transcription, glm.nb() function in MASS, R package (https://cran.r-project.org/web/packages/MASS/index.html; Venables and Ripley 2003), was utilized in enrichment analysis as described in previous research (Supek and Lehner 2017).

### Analysis of mutation rates depending on genomic location

SNVs located in previously described mappable regions were utilized throughout the analysis. Gene information included in “hg19_refGene.txt” file from ANNOVAR tools were utilized to match the additional information of position of SNVs. Classification of sub-genic (5’/3’UTR, protein coding sequence (CDS), and intron) and transcription strand were also based on the gene information. All of the merged genic regions were used as standard for discrimination of genic and intergenic regions. Instead of that, genic regions that do not overlap with other genic regions and CDS regions located between intron regions were utilized for comparing mutation rates in the sub-genic region as previous research reported (Frigola et al. 2017).

### Calling structural variations

Structural variations were called using Delly v0.7.6 (https://github.com/dellytools/delly; Rausch et al. 2012). Raw calls were filtered using in-house scripts. The final call set was manually reviewed by Integrative Genomics Viewer (https://igv.org/; Robinson et al. 2011).

### Calculating RNA expression level

Bulk RNA-seq reads were aligned to GRCh37 using STAR2 v2.6.1d (https://github.com/hbctraining/Intro-to-rnaseq-hpc-O2; Dobin et al. 2013). TPM and reads count were calculated with RSEM v1.3.1 (https://github.com/deweylab/RSEM; Li and Dewey 2011). Differential expression gene analysis was conducted using the DEseq2 package in R (https://bioconductor.org/packages/release/bioc/html/DESeq2.html; Love et al. 2014).

### Calculation of ratio between endogenous APOBEC mRNA and overexpressed APOBEC mRNA

To distinguish transcripts originating from endogenous APOBEC genes versus the exogenous construct, we utilize sequence differences between the two sources. For A3A, two heterozygous variants at chr22:39,357586 (C/T) and 39,357,599 (C/T) were used. In the endogenous A3A allele, reads carried the same base at both positions-either C/C or T/T- while the inserted construct contained C/T bases at the two positions, respectively. For A3B, a homozygous variant at chr22:39,381,999 (C) was present in the endogenous gene, while the construct carried only T at the same locus. The proportion of endogenous mRNA was estimated by calculating the ratio of supporting reads at the positions.

### Calling RNA editings

VarScan2 was utilized to call RNA editing. WGS of mother clones was utilized as paired normal. False positives that did not meet the following criteria were filtered out using an in-house Python script with the pysam module: (1) excluding HLA regions; (2) DNA depth ≥ 5; (3) number of variant reads in DNA = 0; (4) RNA depth ≥ 4; (5) number of variant reads ≥ 3; (6) median number of NM tags in the variant reads < 4; (7) percentage of variant reads with NM tags greater than 3 < 0.5; (8) distance from the end of reads ≥ 5; (9) variants in only one strand; (10) median base quality of variants > 20; (11) percentage of variant reads with clips < 0.5; (12) median mapping quality of variant reads ≥ 30; (13) single-base variant. RNA editings that did not meet all criteria but were commonly observed with all conditions fulfilled in more than two samples were rescued.

To compare the RNA editings, the number of RNA editings was normalized with total sequencing output size. The target output size was determined by the sample having the lowest sequencing output. RNA editings were filtered again by adjusting the ratio of sequencing output to the total depth and variant reads. Variants that met the following two criteria were used in the analysis: (1) corrected depth of RNA ≥ 4 (2) corrected the number of variants.

### Analysis of RNA editing signatures

RNA editing signatures were obtained by a modified version of the mutational signature extraction method used in (Youk et al. 2024; Alexandrov et al. 2013). Briefly, this method uses non-negative matrix factorization (NMF) to disentangle an individual RNA editing spectrum based on a notion of mixed spectra (Cichocki et al. 2006).

RNA editing spectrum was represented as 192 patterns of single-base change with immediately adjacent bases on the 5’ and 3’-end in the canonical mRNA sequence. The number of features is doubled of DNA mutational signature (Alexandrov et al. 2013), because RNA does not necessarily have symmetric base change due to general non-strandedness.

Total 18 experimental data were used as input, split into two subsets: A3A and A3B sets. Each of the A3A and A3B sets included three batches from experiments with 0µg/ml, 0.1 µg/ml, and 3 µg/ml APOBEC3 exposure, totaling 9 batches for each set. Initially, the signature extraction method was used with no additional procedures from the previous method (Youk et al. 2024), and then the optimal number of independent RNA editing signatures was determined. We observed that there were 4 potential candidates for RNA editing signature that are attributable to APOBEC3A, APOBEC3B, ADAR, and the other endogenous editing processes.

APOBEC3A and APOBEC3B aligned well with editing patterns of C>T found in experiments with varying exposure amounts. However, the ADAR signature co-appeared in these signatures at significant proportion. This “leakage” phenomenon occurs due to the constant activity of the ADAR editing process, causing its signal to always appear in RNA editing spectra regardless of the experimental settings. This can be problematic when trying to accurately quantify the attributes of each RNA editing process in a given sample.

To mitigate the potential effect of the leakage phenomenon, an L1 constraint was imposed on the signature matrix (*W*) (Youk et al. 2024). For that, the NMF update algorithm was adopted from a work on sparse NMF (Le Roux et al. 2015). It presents a method that puts L1 constraint on the exposure matrix (*H*). Therefore, we tweaked its objective function by transposing the data matrix (*V*) as well as *W* and *H* and back-transposing them to get *W*. Afterwards, the matrix *W* was normalized so that each column vector sums to one. The sparsity coefficient (μ) was set to 0.6, after trying out several values in a range [0.001, 10].

The A3A set resulted in ADAR and APOBEC3A that were now isolated from each other. The A3B set led to ADAR and APOBEC3B signatures that were also well-separated. ADAR signatures learned from the A3A and A3B set were consistent with cosine similarity 0.999 and were merged into a single ADAR signature by taking average. Endogenously occurring RNA editing other than ADAR was negligible in amount and did not add much value to overall analysis. Thus, we did not seek further signatures.

Lastly, the three RNA editing signatures learned from the data were refitted to individual samples to evaluate the attributes of the RNA editing processes in each sample.

### Analysis of secondary structure of RNA editing sites

Secondary structure of RNA editing sites were predicted as described previously (Jalili et al. 2020). Briefly, a 41 bp sequence centered on each RNA editing site in the canonical mRNA was used to assess the potential for forming secondary structures. Stem strength was calculated as 3*G/C pair + 1*A/T pair in stem. Among candidate structures, the most probable one was selected based on the following hierarchical criteria: (1) highest stem strength, (2) greatest number of G/C pairs in the stem, and (3) smallest loop size.

### Calling clustered mutation

SigProfilerClusters (https://github.com/AlexandrovLab/SigProfilerClusters; Bergstrom et al. 2022a), a python module, was utilized to identify clustered mutations. FlexMix, R package (https://cran.r-project.org/web/packages/flexmix/index.html; Leisch 2004), was utilized to classify the identified clustered mutations into *omikli* and *kataegis*. Since the tools determine the standard intermutation distance for clustering through simulations that randomly distribute SNVs, the total number of SNVs influence the detection rate of clustered mutation events. To correct for the number of clustered mutation events, simulations were conducted.

Identified clustered mutations from our analysis and 703,858 SNVs from 125 samples in PCAWG database, where the contribution of age-related mutational signatures (SBS1, SBS5, and SBS40) is over 0.8, were utilized for simulation. The target number of SNVs was set to 33 (increased by 2,000 from 2,000 to 10,000; increased by 5,000 from 15,000 to 100,000; increased by 10,000 from 110,000 to 200,000). From the observed 615 *omikli* (1,369 SNVs) and 109 *kataegis* (559 SNVs) events, the clustered mutation events were randomly sampled based on the following conditions for 50 iterations: 50, 100, 250, 500 *omikli*, and 25, 50, 75, 100 *kataegis*. Background mutations were selected to match the number remaining after excluding the number of clustered mutations from the target number of SNVs. With the same SigProfilerClusters module, clustered mutations, *omikli and kataegis,* were identified, and the detection rate was calculated. By combining data from the two types of clustered mutations, the relationship between the number of mutations and the detection rate of clustered mutations was estimated (**Supplemental Fig. S13**) using the “drm” function with the fct=AR.3() option in the drc, R package (https://cran.r-project.org/web/packages/drc/index.html; Ritz et al. 2019).

To compare clustered mutation events, 146 samples from the PCAWG database were selected based on the following criteria: (1) nine cancer types showing high prevalence of APOBEC mutational signatures; (2) samples with fewer than 500 SBS2+SBS13 were excluded in both our samples and PCAWG cancer samples to avoid bias.

### Analysis of 4bp context mutations

SNVs found in 10 hGO_iA3A_ samples (excluding control samples) were utilized in the analysis. Among 567 PCAWG samples across eight cancer types with a high prevalence of APOBEC mutational activity - breast adenocarcinoma (BRCA; n=195), esophageal adenocarcinoma (ESAD; n=97), stomach adenocarcinoma (STAD; n=68), head and neck squamous cell carcinoma (HNSC; n=56), lung squamous cell carcinoma (LUSC; n=47), uterine corpus endometrial carcinoma (UCEC; n=44), lung adenocarcinoma (LUAD; n=37), bladder urothelial Carcinoma (BLCA; n=23) - only those with a combined APOBEC-associated mutational burden (SBS2 + SBS13) greater than 5,000 were selected for this analysis (n=63). This subject included: LUSC (n=15), BRCA (n=12), BLCA (n=11), HNSC (n=13), LUAD (n=6), UCEC (n=3), ESAD (n=2), and STAD (n=1).

### Analysis of A3A and A3B expression levels in single cancer cells with public scRNA data

Publicly available single-cell RNA-seq datasets, generated with Smart-seq library, from lung adenocarcinoma, triple negative breast cancer, esophageal adenocarcinoma and esophageal squamous cell carcinoma were analyzed using the same pipeline applied to our bulk RNA-seq data. Epithelial cell populations were first identified following the tutorial workflow provided in the Seurat R package (https://satijalab.org/seurat/; Satija et al. 2015), with the exception of the cell filtering step. Within the epithelial population, cancer cells were distinguished based on the presence of large-scale copy number variations (CNVs) inferred using the inferCNVpy Python package (https://infercnvpy.readthedocs.io/en/latest/index.html). CNV profiles were calculated using fibroblast and endothelial cell populations as reference non-malignant cells. For the head and neck squamous cell carcinoma dataset, where raw data were not available, we utilized a publicly provided summary table containing TPM values and annotated cell types for the analysis.

### Publicly available datasets

Publicly available whole-genome sequencing data were utilized to demonstrate copy number variation in the normal human gastric organoid. WGS of blood from HC05 sample was utilized as the unpaired normal sample (https://ega-archive.org/studies/EGAS00001006213; Nam et al. 2023). To compare the 4bp-context preference and frequency of clustered mutation, SNV calls from the ICGC/TCGA Pan-Cancer Analysis of Whole-Genome (PCAWG) Consortium were utilized for the analysis. The call set data is available for download at https://dcc.icgc.org/releases/PCAWG.

To compare in vivo transcriptional burst of A3A and A3B, we analyzed publicly available Smart-seq-based single-cell RNA-seq datasets from five cancer types: (1) lung adenocarcinoma (https://www.ncbi.nlm.nih.gov/bioproject/591860; Maynard et al. 2020) (2) head and neck squamous cell carcinoma (https://www.ncbi.nlm.nih.gov/bioproject/?term=GSE103322; Puram et al. 2017) (3) triple negative breast cancer (https://www.ncbi.nlm.nih.gov/geo/query/acc.cgi?acc=GSE118390; Karaayvaz et al. 2018) (4) esophageal adenocarcinoma and esophageal squamous cell carcinoma (https://www.ncbi.nlm.nih.gov/bioproject/PRJNA401501; Wu et al. 2018).

### Quantification and statistical analysis

All statistical analyses were performed with R version 4.1.3 (R Core Team, Vienna, Austria). A two-tailed one-sample t-test was used to evaluate p-values for comparing APOBEC-associated SNVs and expression levels between the groups. Linear regressions were conducted using the basic ‘lm’ function in R to analyze the association among APOBEC-associated SNVs, ID9, SBS5, and SBS40. A chi-square test was utilized to evaluate p-values for comparing replication strand bias and transcription strand bias. A 95% confidence interval was used to determine the statistical range of continuous data.

## DATA ACCESS

Whole-genome sequencing, EM-seq, BotSeqS, and bulk RNA-seq raw data produced in this study were deposited in the Korean Nucleotide Archive (KoNA; https://www.kobic.re.kr/kona/). The project accession ID is KAP240815 (https://kbds.re.kr/KRA/browse/Experiment?keyword=KAP240815; https://kbds.re.kr/bioproject_run_all?from_uid=2585828). All data are provided for review purposes upon request by reviewers. Essential in-house scripts used in this study are available on Zenodo (DOI: 10.5281/zenodo.12771074; https://zenodo.org/records/12771074?preview=1&token=eyJhbGciOiJIUzUxMiJ9.eyJpZCI6Ij diNDQ2YzU4LTIxNDMtNDViOS1hYTQ5LTE1MTY4OTEzMmQ2NyIsImRhdGEiOnt9LCJyY W5kb20iOiIwZTg1YmFiYjAzNDRmMzgwOTQ1ZGI2MGI0NDYxMmZkZiJ9.pxeRUXdTXwcoJ rUMkx2ZGlMZNnOyNai2-mcrqyiOIbJkB10lgSOhcvc5Z6mrNKc4TSicAm7NrSgqNops57_SWg).

## ACKNOWLEDGEMENTS

The authors thank Youngwon Cho (Epithelial Biology Center, Vanderbilt University Medical Center) for helping with the cell viability test, Myungsuk Choi (KAIST) for technical help for organoids, and Mia Petljak (New York University) for her valuable advice and feedback on the manuscript. This work was supported by the Young Investigator Grants from the Human Frontier Science Program (RGY0071/2018 to Y.S.J., B.-K.K., and H.S.); the National Research Foundation of Korea (Leading Researcher Program NRF-2020R1A3B2078973 to Y.S.J.); the Korea Bio Data Station (N24NM016-24 to J.L and J.W.P); and the Institute for Basic Science (IBS-R021-D1 to B.-K.K).

## Author Contributions

Y.S.J., B.-K.K., and H.S. designed the experiments. J.-H.K. provided human gastric epithelium sample from biopsy. Y.J.B. established organoids from primary human gastric tissues. J.L. and J.W.P. helped basic bioinformatic works (alignment and calling somatic variants). J-H.L. and J.Y. conducted vector cloning. Y.A. and T.K. conducted the transformation of organoids, with J-H.L. and J.Y. providing advice. Y.A. performed organoid culture, clonal expansion, and DNA/RNA extraction. B.-K.K. and H.K. provided training in organoids culture technologies. Y.A. performed cell viability analysis. J-H.L. performed immunohistochemistry. S.A.O. conducted *in vitro* deamination experiments and most of the library construction including WGS and RNA-seq. Y.A., J.-H.P., and K.Y. conducted library construction for duplex sequencing analysis. Y.A. conducted most bioinformatic analyses, including alignment, mutation calling, duplex sequencing analysis, and analysis of expression levels with bulk RNA-seq data, with S.P., J.Y., and Y.S.J. providing advice. Y.A. and Jo.L. conducted quantitative and statistical data analysis. Y.A. and Jo.L. conducted mutational signature analysis including RNA editing. W.H.L. helped with the construction of pipeline for analysis of duplex sequencing. Y.A. and C.H.N. conducted epigenomic analysis of variants.

