## Supplemental Fig. S1 for "APOBEC3A, not APOBEC3B, drives deaminase mutagenesis in human gastric epithelium"

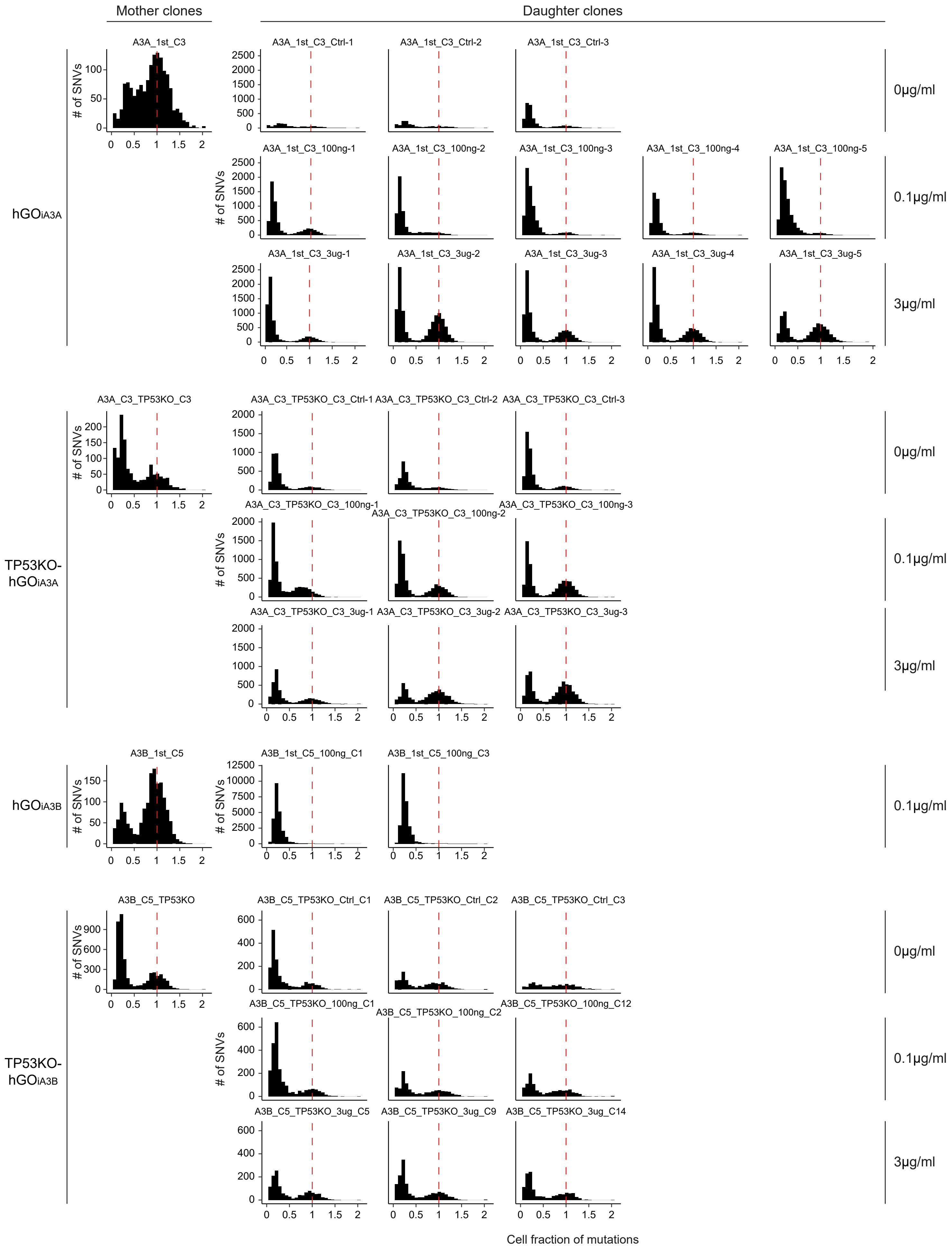


**Supplemental Figure S1. Distribution of cell fraction of mutations in each clone used in this study.** Each cell fraction of mutation represents the proportion of cells harboring each single nucleotide variant.
