## Supplemental Fig. S2 for "APOBEC3A, not APOBEC3B, drives deaminase mutagenesis in human gastric epithelium"

**A**

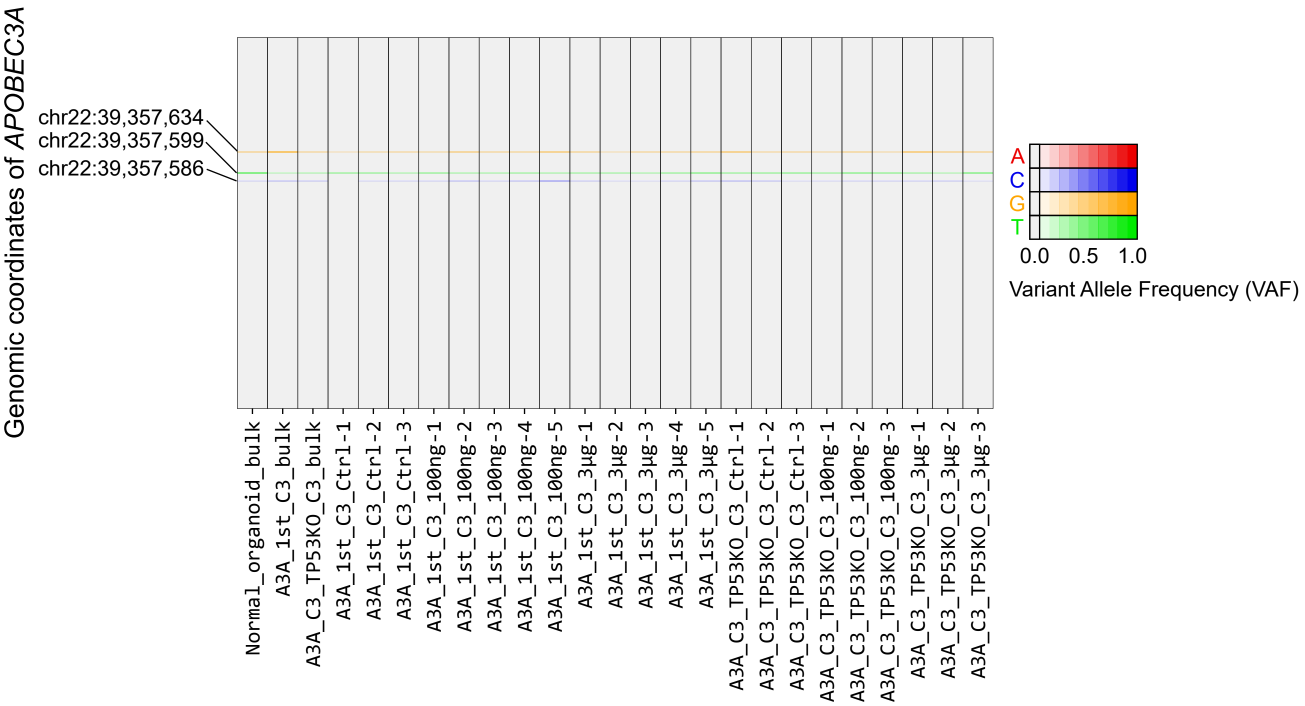


**B**

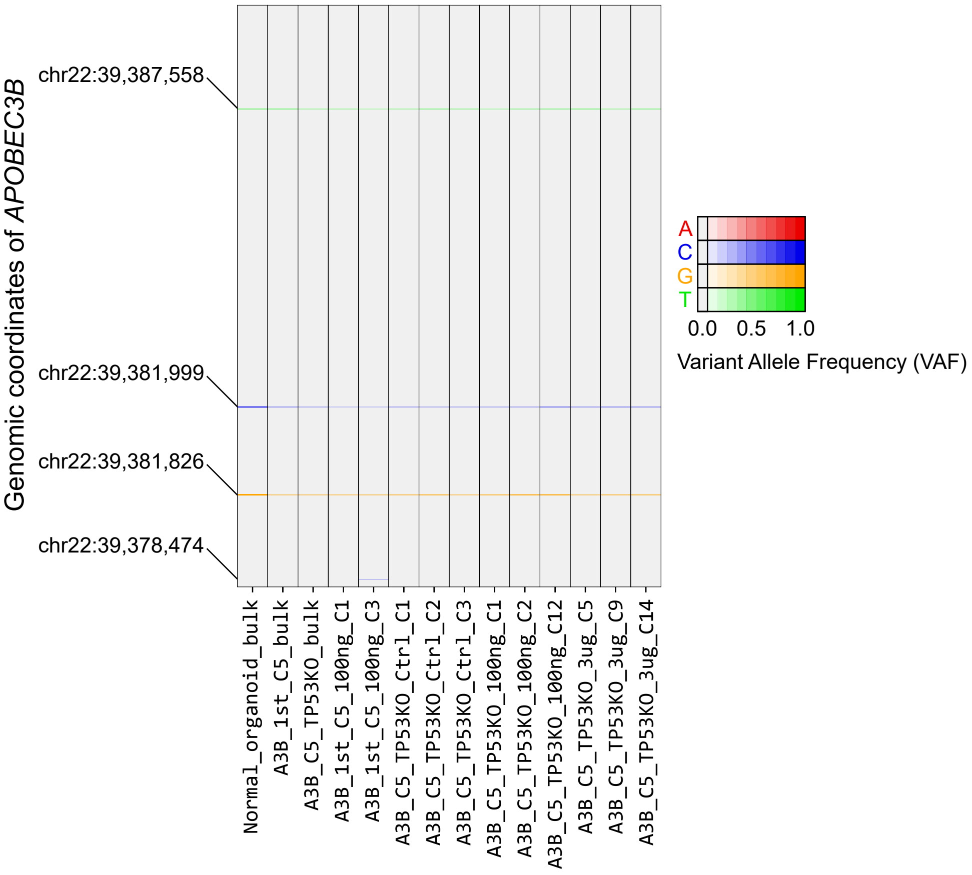


**Supplemental Figure S2. Variant allele frequencies (VAFs) against reference APOBEC sequence in each clonal organoid line. (A)** hGOiA3A and TP53KO-hGOiA3A **(B)** hGOiA3B and TP53KO-hGOiA3B. All missense mutations annotated with genomic positions originated from either the *A3A* or *A3B* in endogenous copies, except for one missense mutation (chr22:39,378,474) in A3B_1st_C5_100ng_C3 sample, which resulted from a sequencing error.
