## Supplemental Fig. S3 for "APOBEC3A, not APOBEC3B, drives deaminase mutagenesis in human gastric epithelium"

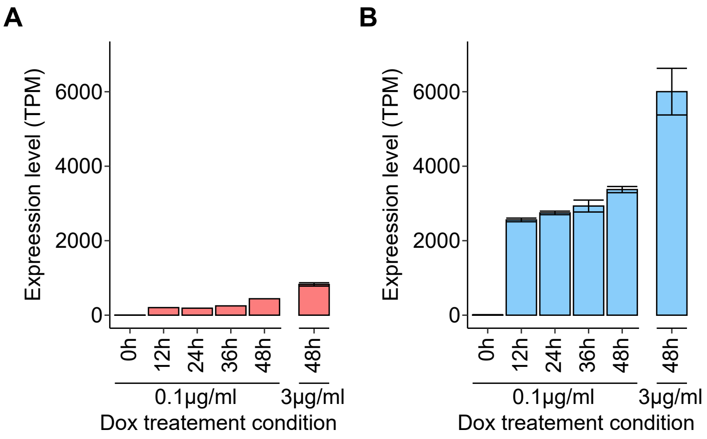


**Supplemental Figure S3. Expression levels of A3A or A3B in each corresponding organoid line** **(A)** Expression levels of APOBEC3A (A3A) in $hGO_{iA3A}$ line following under each doxycycline condition (n=3 in each condition). **(B)** Expression levels of APOBEC3B (A3B) in $hGO_{iA3B}$ line following under each doxycycline condition (n=3 in each condition). Data are presented as mean ± 95% confidence intervals.
