## Supplemental Fig. S4 for "APOBEC3A, not APOBEC3B, drives deaminase mutagenesis in human gastric epithelium"

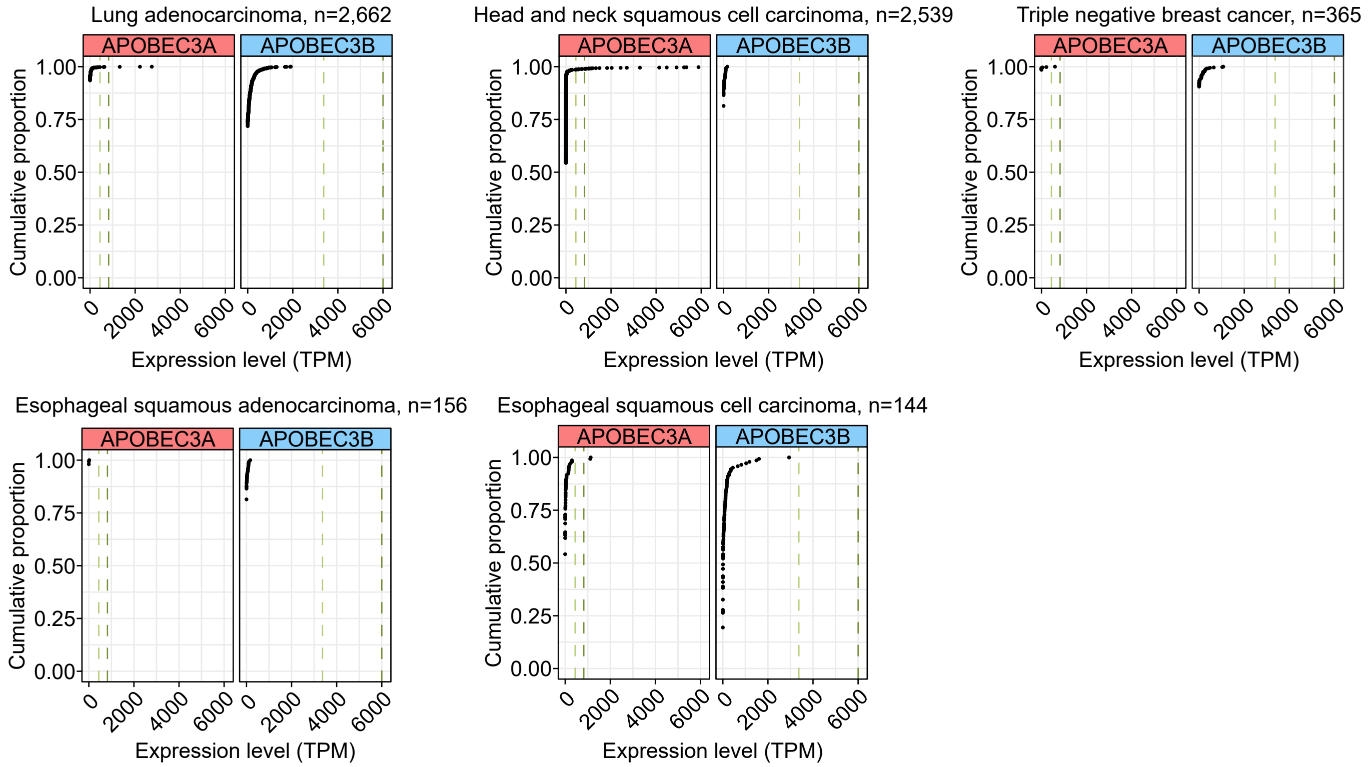


**Supplemental Figure S4. Cumulative proportions of expression levels of A3A and A3B in single cancer cells across multiple types of cancer.** Green line; average expression levels of A3A and A3B following 0.1 µg/ml doxycycline for 48 hours in each corresponding line, $hGO_{iA3A}$ and $hGO_{iA3B}$, respectively; dark green line; average expression levels of A3A and A3B following 3 µg/ml doxycycline for 48 hours in each corresponding line, $hGO_{iA3A}$ and $hGO_{iA3B}$, respectively.
