## Supplemental Fig. S5 for "APOBEC3A, not APOBEC3B, drives deaminase mutagenesis in human gastric epithelium"

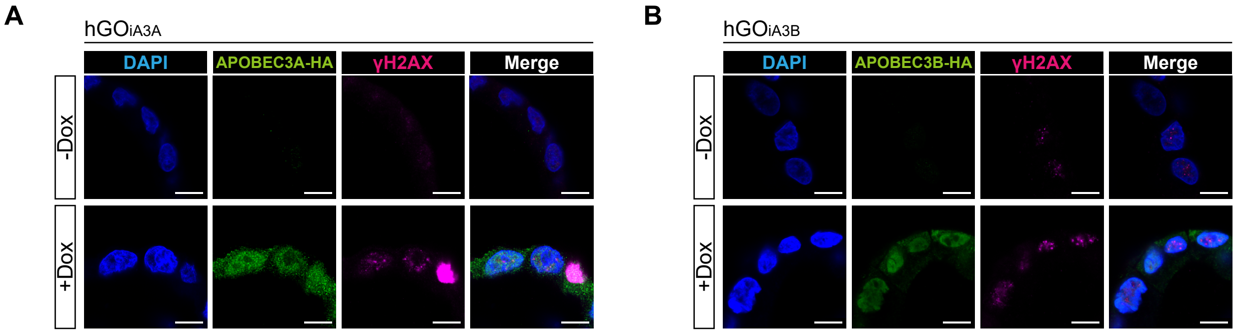


**Supplemental Figure S5. High resolution images of immunohistochemistry. (A)** hGO$_{iA3A}$ and **(B)** hGO$_{iA3B}$ lines. Scale bars represent 10μm.
