## Supplemental Fig. S6 for "APOBEC3A, not APOBEC3B, drives deaminase mutagenesis in human gastric epithelium"

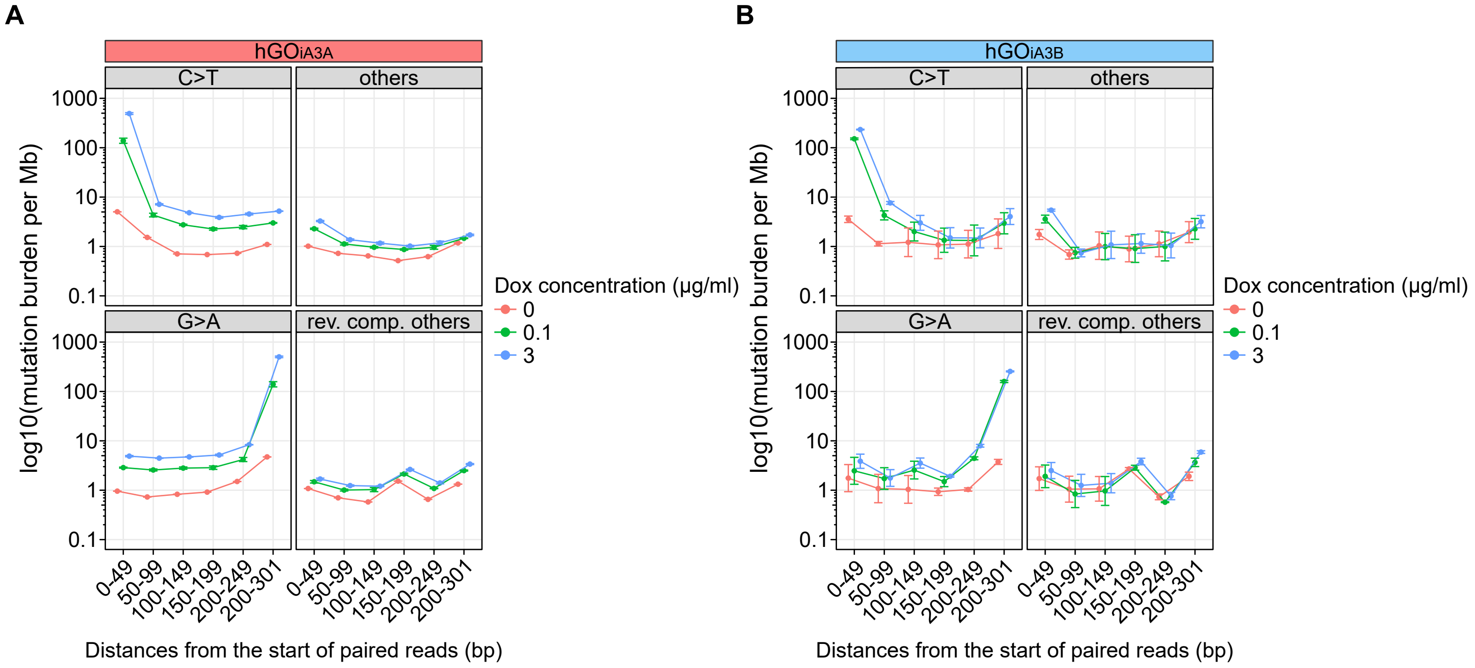


**Supplemental Figure S6. Distribution of distances from the start position of paired reads to each single nucleotide variant (SNV) in BotSeqS results in each line. (A)** hGOiA3A and **(B)** hGOiA3B lines. The 0-position is the 5' head region of each DNA fragment.
