## Supplemental Fig. S7 for "APOBEC3A, not APOBEC3B, drives deaminase mutagenesis in human gastric epithelium"

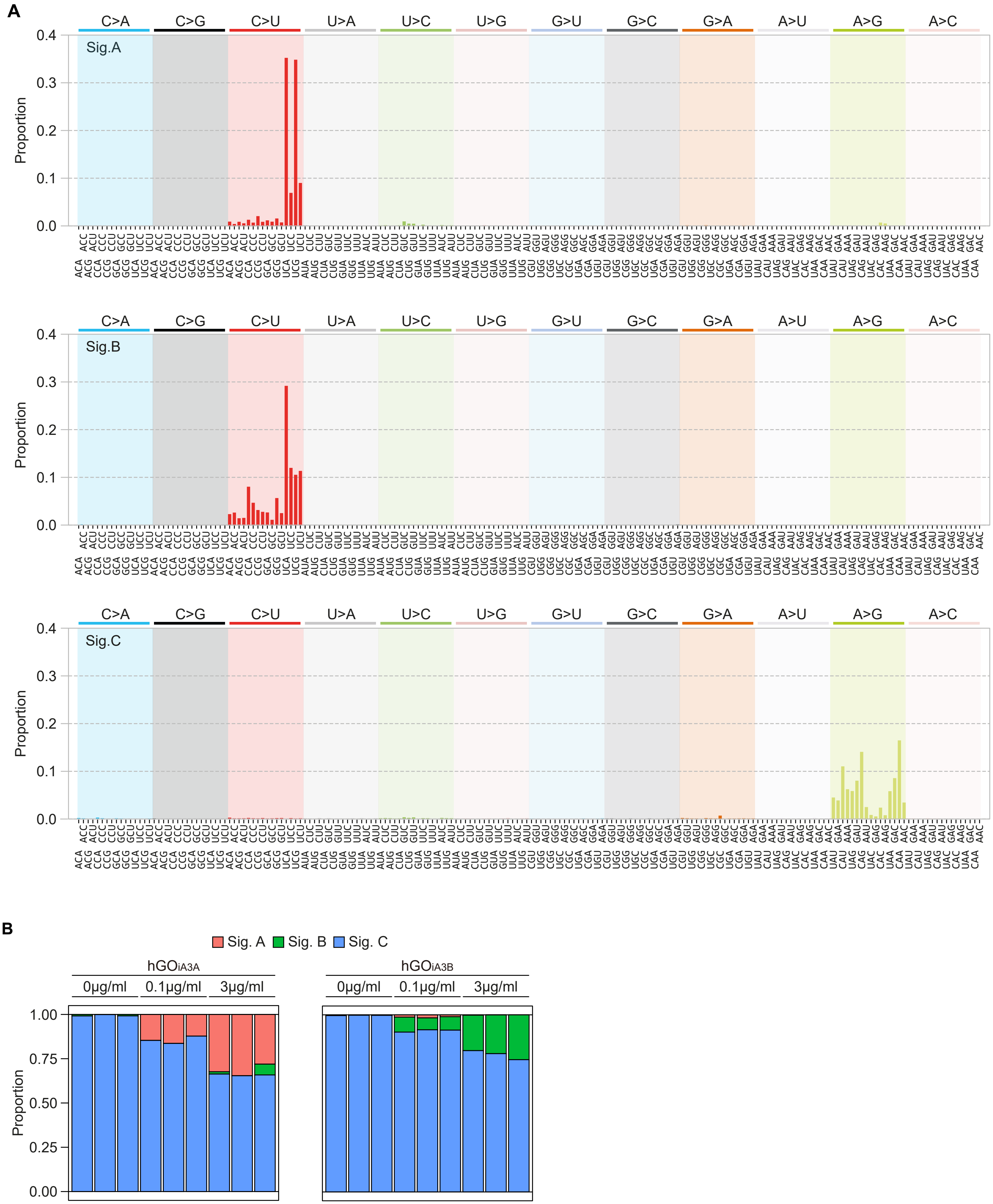


**Supplemental Figure S7. RNA editing signatures in the hGO$_{iA3A}$ and hGO$_{iA3B}$ clones. (A)** Decomposed spectra of RNA editing signatures across doxycycline treated hGO$_{iA3A}$ and hGO$_{iA3B}$ lines. **(B)** Proportion of each decomposed RNA editing signature in each hGO$_{iA3A}$ and hGO$_{iA3B}$ line under the doxycycline treatment conditions.
