## Supplemental Fig. S8 for "APOBEC3A, not APOBEC3B, drives deaminase mutagenesis in human gastric epithelium"

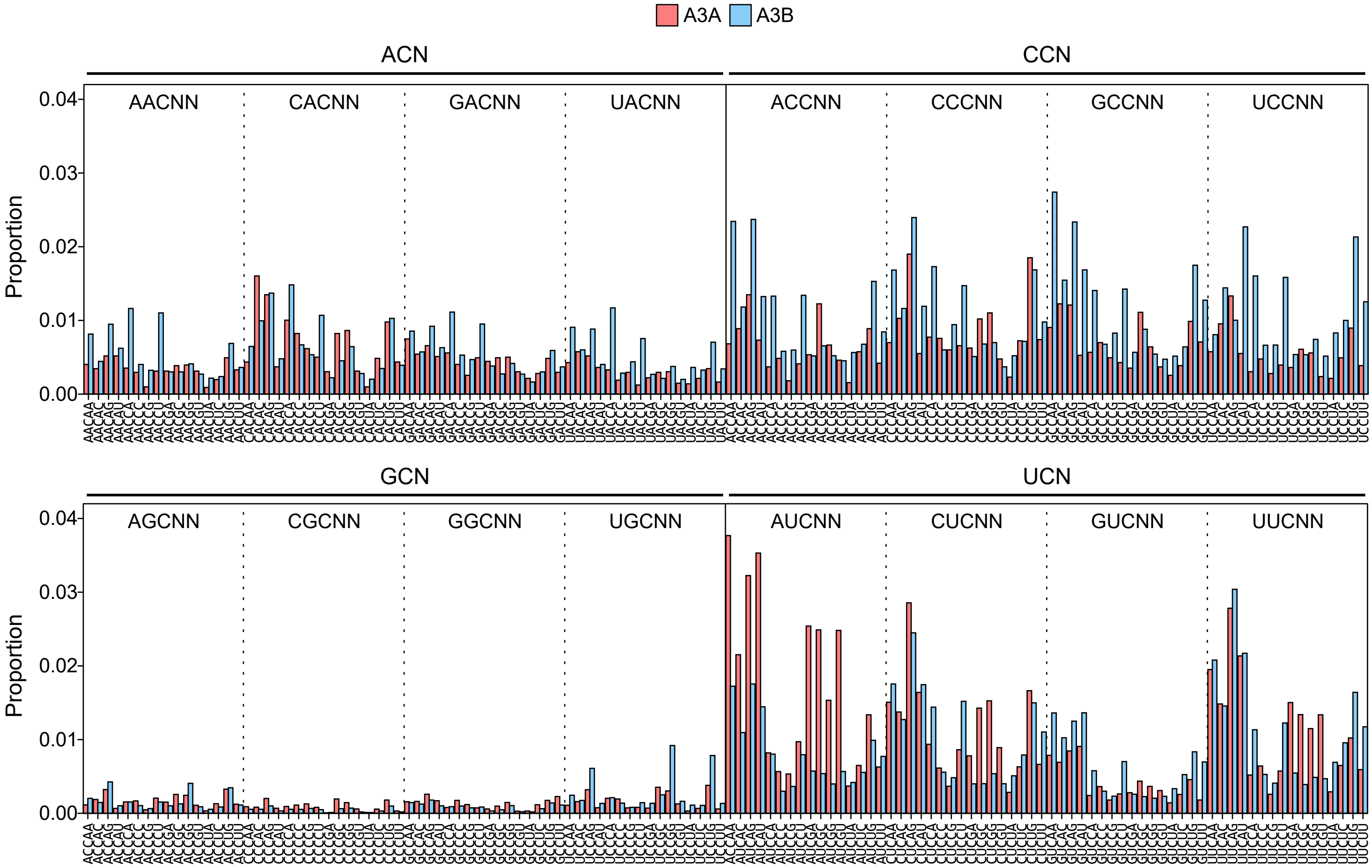


**Supplemental Figure S8. Spectra of C>U RNA editing in pentanucleotide contexts from hGO$_{iA3A}$ and hGO$_{iA3B}$ lines after 3μg/ml doxycycline treatment for 48 hours.**
