## Supplemental Fig. S9 for "APOBEC3A, not APOBEC3B, drives deaminase mutagenesis in human gastric epithelium"

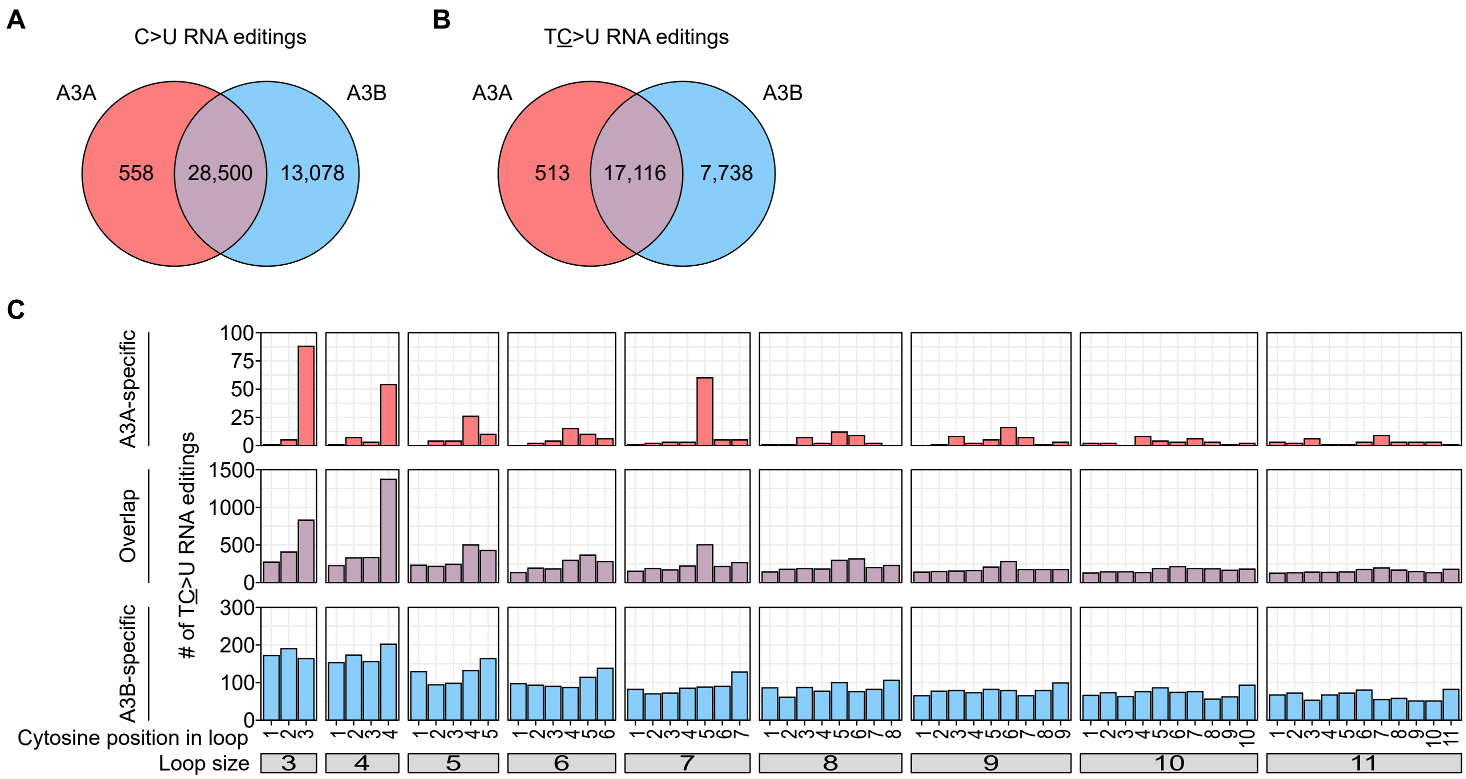


**Supplemental Figure S9. Characteristics of APOBEC-associated C>U RNA editing recurrent sites in the hGO$_{iA3A}$ and hGO$_{iA3B}$ clones.** RNA-seq results after doxycycline treatment for 48 hours with 0µg/ml, 0.1µg/ml, and 3µg/ml were utilized for the analysis. **(A)** Venn diagram showing the number of C>U RNA editing recurrent sites (n ≥2 in each organoid line). **(B)** Venn diagram showing the number of TC>U RNA editing recurrent sites. **(C)** Properties of the proposed secondary structures of the TC>U RNA editing recurrent sites.
