## Supplemental Fig. S10 for "APOBEC3A, not APOBEC3B, drives deaminase mutagenesis in human gastric epithelium"

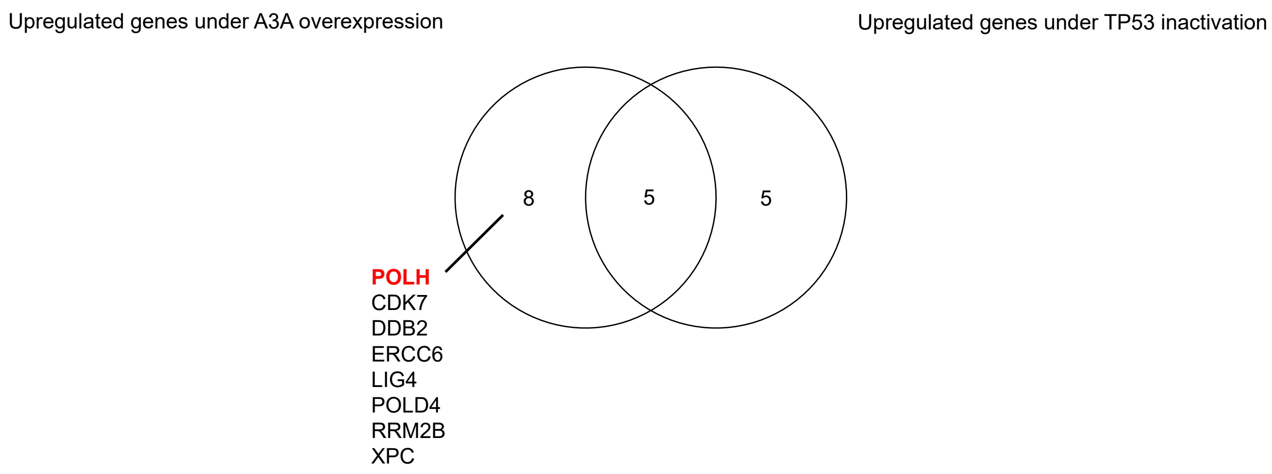


**Supplemental Figure S10. Differentially expressed genes contributing DNA repair in hGO$_{iA3A}$ and TP53KO-hGO$_{iA3A}$ lines under A3A overexpression with 3μg/ml doxycycline treatment for 48 hours.** The numbers in the Venn diagram represent the number of differentially expressed genes belonging to each comparison group.
