## Supplemental Fig. S11 for "APOBEC3A, not APOBEC3B, drives deaminase mutagenesis in human gastric epithelium"

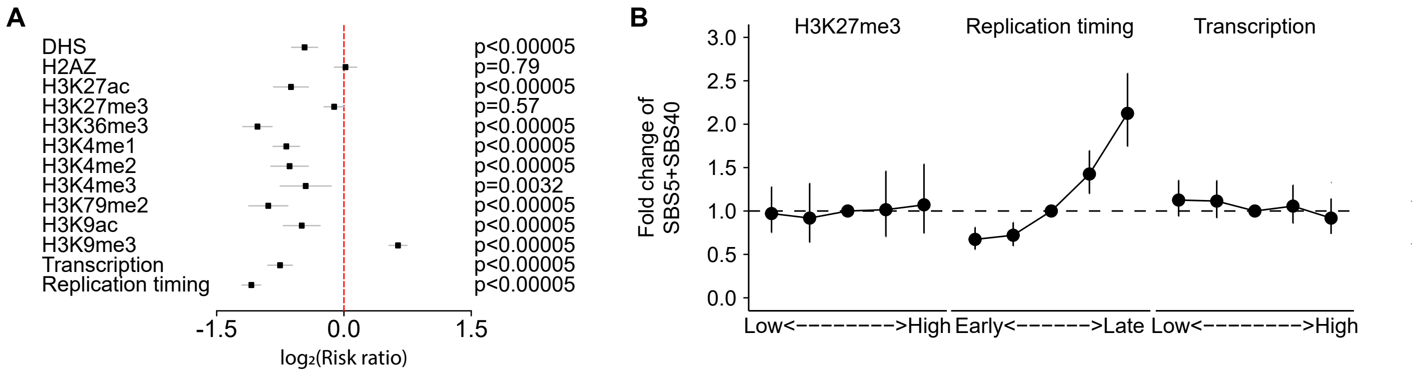


**Supplemental Figure S11. Genomic and epigenomic distribution of mutations contributing SBS5 and SBS40. (A)** Correlation between epigenetic markers and SBS5- and SBS40-associated substitutions. **(B)** Fold change of mutation rates of SBS5- and SBS40-associated SNVs according to the replication.
