## Supplementary figures and images for "APOBEC3A, not APOBEC3B, drives deaminase mutagenesis in human gastric epithelium"

### Supplemental Fig. S12

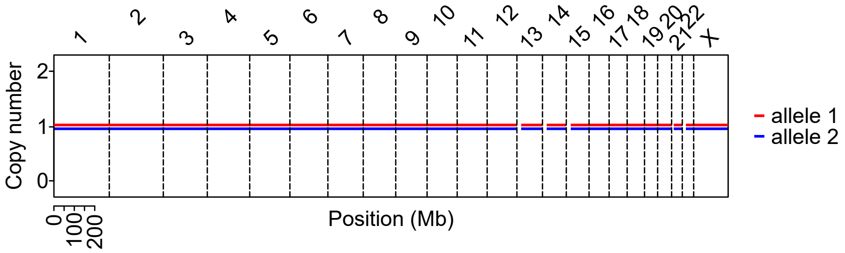


**Supplemental Figure S12. Copy number variations of normal gastric organoids.**
