## Supplemental Fig. S13 for "APOBEC3A, not APOBEC3B, drives deaminase mutagenesis in human gastric epithelium"

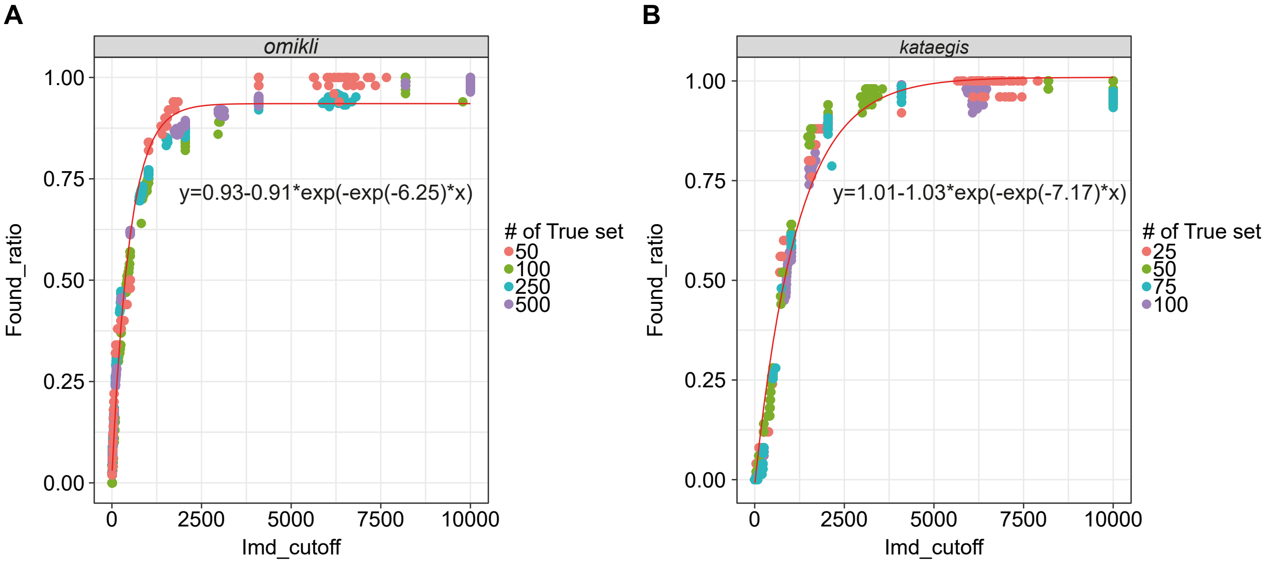


**Supplemental Figure S13. Detection rate of clustered mutation depending on total number of mutations. (A)** Detection rate of *omikli* depending on the total number of mutations. **(B)** Detection rate of *kataegis* depending on the total number of mutations.
